# MAM-STAT3-induced upregulation of mitochondrial Ca^+2^ causes immunosenescence in patients with type A mandibuloacral dysplasia

**DOI:** 10.1101/2022.08.31.504639

**Authors:** Arshad Ahmed Padhiar, Xiaohong Yang, Zhu Li, Jinqi Liao, Ilyas Ali, Wei Shu, AA Chishti, Liangge He, Gulzar Alam, Abdullah Faqeer, Yan Zhou, Shuai Zhang, Ting Wang, Tao Liu, Meiling Zhou, Gang Wang, Xuenong Zou, Guangqian Zhou

## Abstract

Homozygous lamina/c p.R527C mutations result in severe mandibuloacral dysplasia (MAD) and progeroid syndrome, but the underlying molecular pathology remains unknown. Here, we report on three patients with MAD, all displaying severe systemic inflammaging and characterized the major molecular pathways involved in the manifestation of this disease. Analysis of induced pluripotent stem cell (IPSC)-derived mesenchymal stem cells (MAD-iMSCs) obtained from the patients revealed that increased mitochondrial Ca^+2^ loading was the root cause of lost mitochondrial membrane potential, abnormal fission/fusion and fragmentation, which then participated in inflammaging by inducing the inflammasome. These alterations in Ca^+2^ homeostasis were mediated by signal transducer and activator of transcription 3 (STAT3), which is located on the mitochondrial associated membrane (MAM). STAT3 function could be rescued by treatment with clinically-approved IL-6 blockers, or by correction of R527C mutations. In addition, extracellular vesicles (EVs) obtained from MAD-iMSCs displayed reduced immunomodulatory function, being unable to rescue bleomycin-induced lung fibrosis and triggering mitochondrial dysfunction, senescence, and fibrosis in healthy cells. Our results provide new insights into the pathology of complex lamin-associated MAD with systemic immunosenescence, and suggest that targeting defective mitochondrial Ca^+2^ homeostasis may represent a promising novel therapy for this condition.

## INTRODUCTION

Laminopathies are a spectrum of rare, degenerative genetic disorders involving defects in the nuclear lamina, and are characterized by accelerated aging. These conditions are primarily associated with mutations in the lamin A/C (LMNA) gene, which codes for both lamin-A and lamin-C proteins via alternative splicing. Lamin-A and lamin-C serve as integral components of the nuclear lamina, where they provide mechanical support and ensure genomic stability^1^. To our knowledge, over 500 different LMNA mutations have been linked to range of diseases, including lipodystrophies, cardiac myopathy and muscular dystrophy, as well as the severe, systemic laminopathy known as Hutchinson-Gilford progeria syndrome (HGPS)^2^. Around 90% of HGPS cases are caused by a de novo LMNA mutation (c.C1824C>T; G608G) which activates aberrant mRNA splicing and the subsequent truncation and permanent farnesylation of lamin-A (also known as progerin)^1^. Accumulation of progerin culminates in premature senescence, leaving patients with HGPS with an average life expectancy of 13 years^3^. However, other homozygous and compound heterozygous point mutations in LMNA have been found to cause similar pathologies. These conditions are commonly referred to as mandibuloacral dysplasia (MAD) or mandibuloacral dysplasia with type-A lipodystrophy (MADA), and are characterized by acro-osteolysis, mandibular hypoplasia, lipodystrophy and varying degrees of accelerated aging^4^.

MADA is vague medical term, and does not always reflect the severity of the disease or its time of onset. For example, patients bearing homozygous LMNA p.R527H mutations typically begin to experience mild MAD symptoms during their second decade of life^5^. However, in patients with homozygous p.R527C mutations, symptoms arise as early as 10 months and become severe by the age of 4 years^6^. These differences were explained by a computational model, which predicted that certain types of mutation in the highly-conserved lamin a/c immunoglobulin (Ig)-like domain disrupt the nuclear envelop more than other mutations^7^. Furthermore, the nuclear envelope proteome is highly variable between different tissues, which can cause disease to manifest in a tissue-specific manner in people with LMNA mutations. Thus, drugs approved to treat HGPS, such as lonafarnib^8^, would likely fail to treat MAD or atypical progeria. As these disorders are so rare, the most practical approach would be to identify underlying common pathological features that are involved in MADA and attempt to target these. Notably, a recent study has reported that biallelic mutations in the outer mitochondrial protein metaxin-2 (MTX-2) also result in severe MAD, with atypical progeroid symptoms^9^. This suggests that progeria can also arise in the absence of primary defects in the nuclear lamina network. A deeper understanding of progeroid pathology in the context of mitochondrial defects might aid in the development of treatments for a range of laminopathies.

Universal Mutation Database (UMD-LMNA; http://www.umd.be/LMNA/) archived ∼3000 subjects, associated with 500 different LMNA mutation. Compared to others, homozygous LMNA pR527C manifest towards systemic aging with faster pace and presents natural model to comprehend other aging-associated diseases. The number of MAD cases bearing homozygous LMNA p.R527C mutation has increased in Southern China in recent years^4, 10^, suggesting the founder effect. In this study, we report on three new cases of MAD in patients with LMNA p.R527C mutations, and provide the first characterization of the underlying pathology. Our results suggest that deregulated mitochondrial calcium signaling is the primary cause of the inflammaging and premature senesce associated with MAD disorders. It may be possible to treat both typical and atypical laminopathies by targeting the mechanisms underlying these mitochondrial defects with currently-approved drugs, which would facilitate rapid adoption in clinical settings.

## Results

### Homozygous LaminA/C p.R257C patients present with severe MAD and immunosenescence

A total of four children from three distinct ethnic groups were admitted to hospital with overlapping progeroid symptoms, including a large head, sparse hairs, a pinched nose, a high-pitched voice, and subcutaneous lipoatrophy. We initially conducted full-length sequencing of the common genes involved in progeroid syndromes, including LaminA/C, ZMPSTE24, POLD1 and BNF1. Sequencing of these genes revealed that three patients had homozygous base pair mutations in exon 9 of LaminA/C c.1579C/T (p.R527C). These patients were subsequently referred to as MAD1 (female, 3yr), MAD2 (male, 5yr) and MAD3 (male, 7yr). The fourth child (male, 1yr) bore a heterozygous c.1824C/T (p.G608G) mutation of LaminA/C, which is associated with the development of typical progeria. This final patient was subsequently referred to as HGPS1. The parents and siblings of patients MAD1-3 were heterozygous and asymptomatic for mutation (Fig. 1A and B), suggesting a pattern of autosomal recessive inheritance. However, no mutation was observed in the parents of patient HGPS1, suggesting that G608G represents a sporadic dominant mutation. Detailed clinical features, including radiological observations, are described in Supplementary information, Note 1 and Fig. S1.

**Fig. 1.**
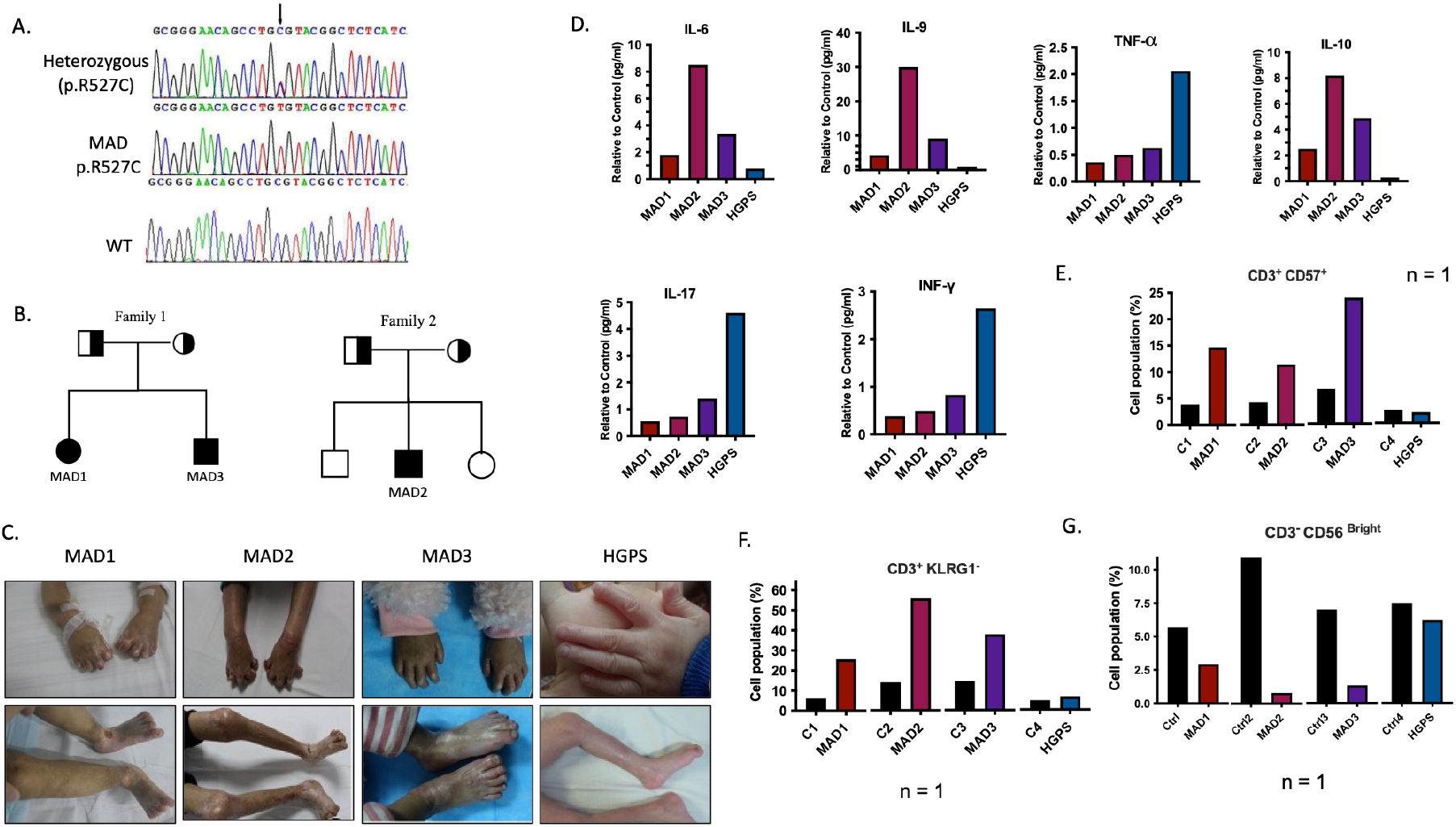
Clinical and molecular pathological features of LMNA p.R527C MAD patients. (A) Example electropherograms for a patient bearing homozygous p.R527C mutations, their asymptomatic, heterozygous siblings/parents, and the wild-type healthy control. (B) The pedigrees of two ethnically-distinct families with LaminA/C p.R527 mutations. Squares represent males, circles represent females; and filled, half-filled and unfilled symbols reflect affected individuals, asymptomatic carriers and non-carriers of the mutations, respectively. (C) Images of patients with homozygous LaminA/C p.R527C mutations showing mandible hypoplasia with crowded teeth, severe contracture at the interphalangeal joints, flexion deformity of the fingers, club-shaped phalanges or rounding of fingertips with marked acro-osteolysis. Frequent ulceration and scleroderma was also observed on the lower trunk of patients with MAD. Reduced scalp hair, decreased eyebrows, beaked nose, fullness of cheeks and skin thickening around the ankle with stiff joints were common between all patients. (D) Cytokine levels were quantified from patient serum and expressed as fold change after normalizing with healthy, age-matched controls. (E-G) Data are presented as relative values. Patients: MAD1 (Female, 3 y); MAD2 (Male, 5 y), MAD3 (Male, 7 y). Controls: C1 (Female, 3 y); C2 (Male, 5 y); C3 (Male, 7 y); C4 (Male, 1 y).

The three homozygous LMNA R527C patients presented with severe MAD symptoms, which resembled those of auto-immune disorders such as rheumatoid arthritis. (Fig. 1C). Examination of skin biopsies revealed the infiltration of immune cells and severe dermal sclerosis (Supplementary information, Fig. S2). However, the serum antibodies for antinuclear and/or anti-smith (sm) were below the reference values (Supplementary information, Table S1), suggesting that the overactivation of the immune response was not caused by an auto-immune disorder in these patients.

Polarization of immune responses into either inflammatory or anti-inflammatory responses is based on the cytokines released by T-helper (TH) cells. Thus, we profiled different TH mediated cytokine levels, and found that IL-6, IL-9 and IL-10 expression was higher in serum taken from patients MAD1-3 compared with patient HGPS1, while IL-17F, TNF-α and INF-γ levels were higher in patient HGPS1 compared with patients MAD1-3 (Fig. 1D; Supplementary information, Table S2). Furthermore, the T lymphocyte, B cell and NK cell populations of patients did not differ from those of controls, although increased levels of terminally-differentiated senescent T and NK lymphocyte markers were detected in MAD1-3 patients (Fig. 1E-G; Supplementary information, Table S3-S6).

Unlike in the patient serum, the levels of cytokines from cultured MAD peripheral blood mononuclear cells (PBMCs) were not significantly different than those found in the controls (Supplementary information, Table S7 and S8). This implies that other cells or organs are involved in mediating immunosenescence in these patients. Taken together, these results suggested the hyperactive immune system observed in patients with MAD and HGPS occurs via different patterns of cytokine expression.

### MAD patient-derived iMSCs represent a suitable model for studying premature senescence

The availability of physiologically-relevant models is a crucial step in disease etiology and drug discovery. Induced pluripotent stem cells (IPSCs) can faithfully recapitulate both the clinical manifestation and pathophysiology of complex multi trait disorders, and so serve as suitable models. To this end, PBMCs from patients with MAD were reprogrammed using Yamanaka factors (Klf4–Oct3/4–Sox2 and c-myc), and were termed IPSC-MAD1, IPSC-MAD2 and IPSC-MAD3. From among several positive IPSC-like colonies, three clones from each patient with MAD were selected after confirming a normal karyotype, the presence of embryonic stem cell markers and the ability to differentiate into three embryonic germ layers via teratoma formation (Supplementary information, Fig. S3 and S4). The expression of LaminA/C in these IPSCs was sparse, albeit present (Fig. 2A; Supplementary information, Fig. S5A, B), which contracted previous studies that showed a complete attenuation of LaminA/C expression at both the RNA and protein level^11, 12^. This discrepancy could be due to changes in the starting cell population or variations in the reprogramming methodology. However, the pluripotent markers were maintained after >40 passages (Supplementary information, Fig. S5C), indicating successful reprograming from the premature senescent cell phenotype.

**Fig 2.**
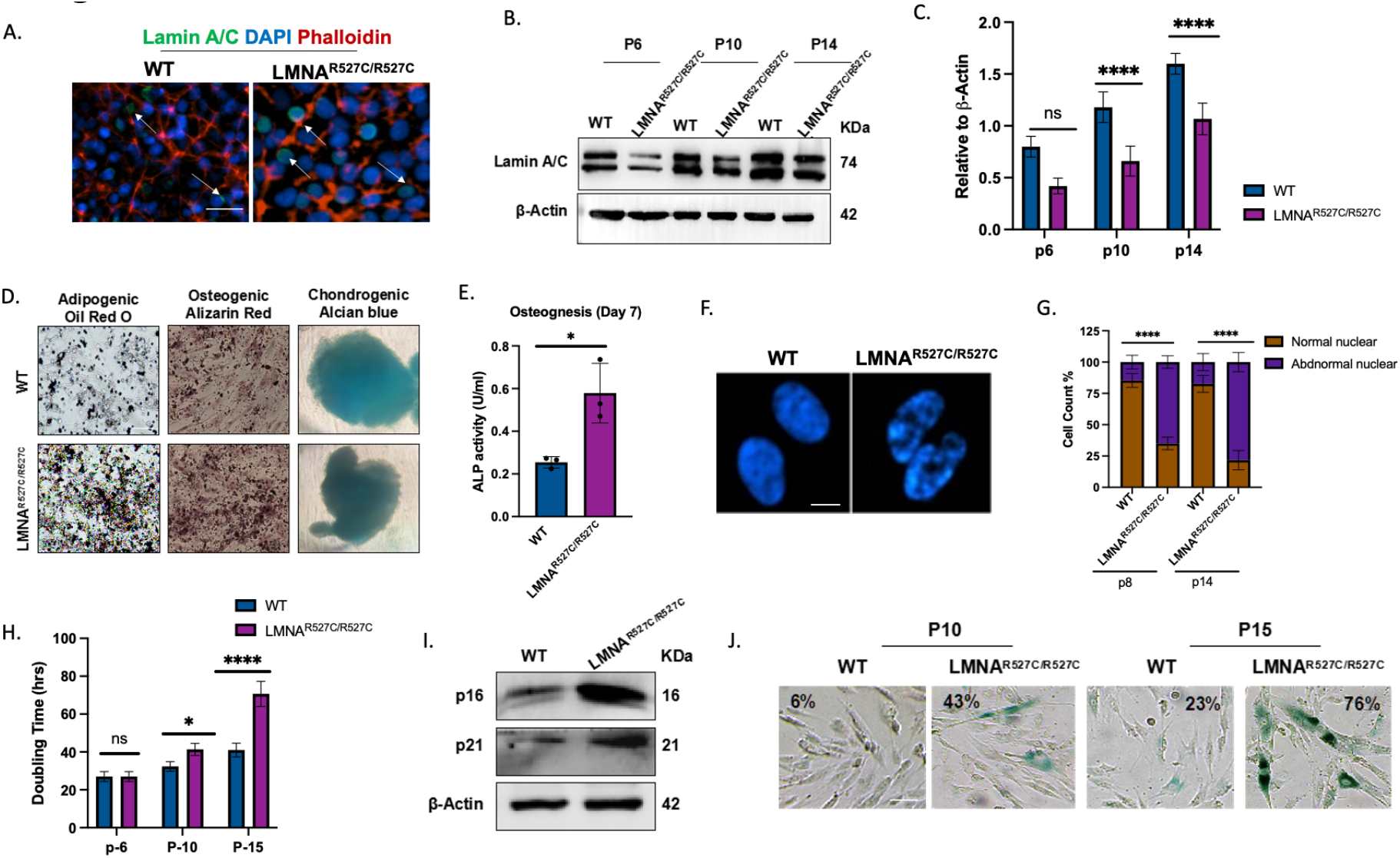
MAD iMSCs showed accelerated senescence and impaired Adipogenesis and osteogenesis. In all the cases the error bar represents standard deviation. *p<0.05, **p<0.01, and ***p<0.001, with comparisons indicated by lines. (A) Indirect Immunofluorescence staining on IPSCs (passage 22) against LaminA/C (green). Nucleus stains with DAPI (blue) and red showed counter staining using phalloidin. The scale bar represents 100 µm. (B and C) Western blot analysis showed that laminA/C expression increased with passage number, although this expression was significantly lower in LMNA^R527C/R527C^ iMSCs compared with WT-iMSCs. (D) Tri-lineage differentiation of iMSCs (n=3). iMSCs were differentiated for 18 days and then stained with oil-red-oil dye, which showed the accumulation of more lipid droplets in MAD-iMSCs than control. Alizarin staining was used to assess osteogenic differentiation after 14 days of induction. MAD-iMSCs displayed higher nodular structure with more calcium content compared to wild type iMSCs. Alcian blue dye showed no difference in MAD-iMSCs with control in their ability to form cartilage matrix, rich in aggrecan. Aggrecan was stained after 21 days of differentiation (n = 5). Scale bar represent 1000 µm. (E) Alkaline phosphatase enzyme levels were measured in iMSCs on day 7 of differentiation into osteogenesis (n =3). (F and G) Fluorescence images of nuclei stained with DAPI showed abnormal nuclear morphology in MAD-iMSCs, with nuclear blebbing and honeycomb nuclei. The scale bar represents 20 µm (n=8). (H) Cell proliferation plotted against cell doubling time (n =3). (I) Western blot analysis of whole cell lysates at passage 10. (J) Cell senescence was quantified at the indicated passage in MAD-iMSCs and WT-iMSCs with beta-galactosidase staining. Scale bar represents 200 µm.

**Fig 3.**
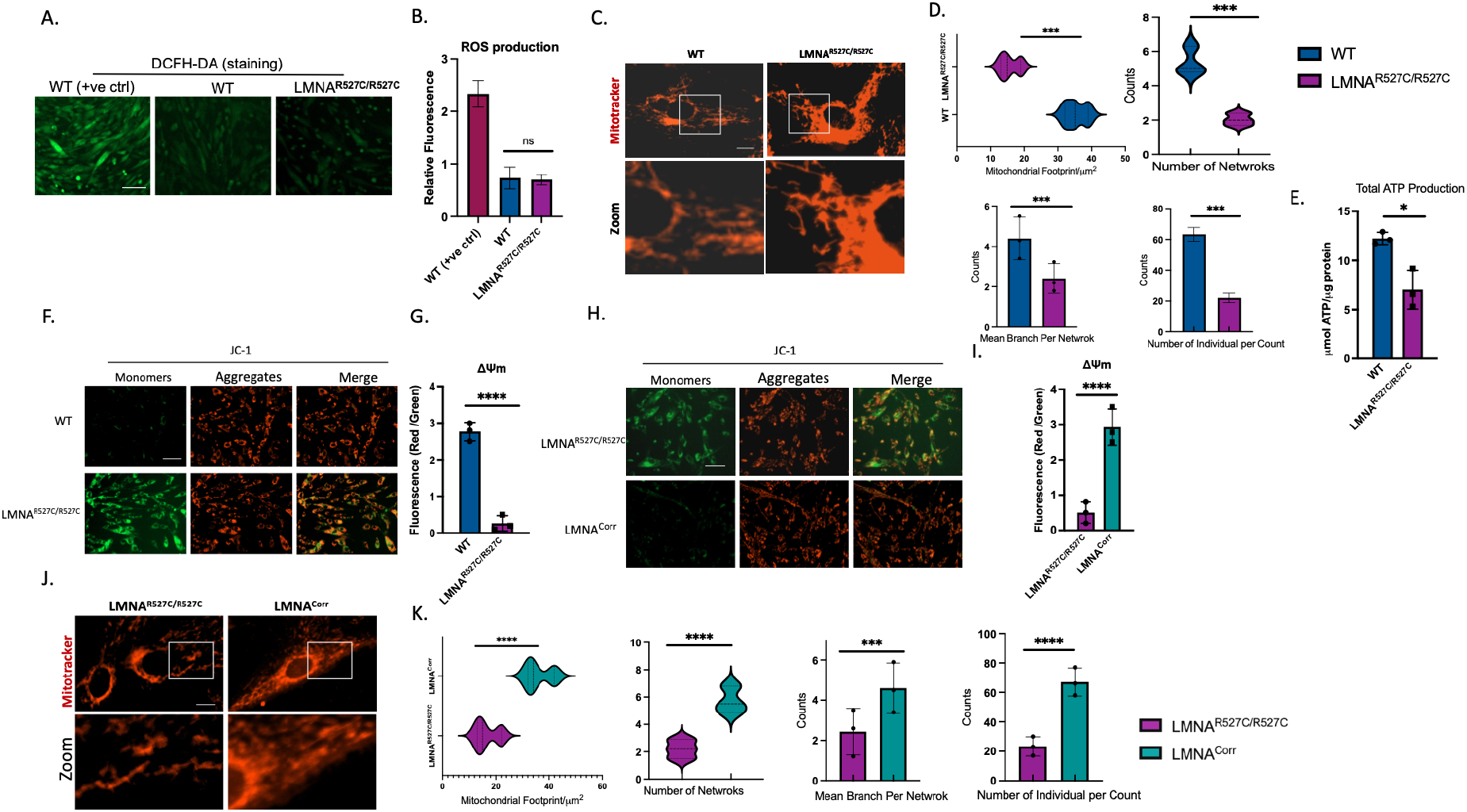
Mitochondrial dysfunction in MAD patient iMSCs is rescued by CRISPR/CAS9 correction. All experiments present in the figure conducted between 7 to 10 cell passage number. *p<0.05, **p<0.01, and ***p<0.001, with comparisons indicated by lines. (n = ≥ 4). (A and B) DCFH-DA staining revealed no significant change in MAD or wildtype iMSCs. Healthy iMSCs were treated separately with Rosup, which induces oxidative stress, and were used as a positive control. Scale bar = 200 µm. (C and D) Representative immunofluorescent staining revealed the mitochondrial network morphology by staining with Mitotracker (red), scale bar represent 20 µm. There was a significant difference in mean branch per network, number of individual/count, mitochondrial foot prints and total number of mitochondrial networks in MAD-iMSCs compared with with wild type cells. (E) ATP production was quantified and presented as relative to total protein concentration. (F and G) Representative images of mitochondrial membrane potential, quantified using JC-1 dye. Scale bar = 200 µm. (H and I) Mitochondrial fragmentation was rescued when MAD-iMSCs was corrected with CRISPR/CAS9, as described in method section. Scale bar = 200 µm. (K and L) Mitochondrial membrane potential, which was analysed with JC-1 dye, showed less monomer (green) in CRISPR/CAS9 corrected iMSCs. Scale bar = 20 µm

Senescent mesenchymal stem cells (MSCs) serve a vital role in immunosenescence, which leads to the chronic subclinical inflammatory state known as inflammaging. Therefore, we hypothesized that MSCs may represent an ideal model for understanding the mechanisms underlying the inflammaging observed in LMNA R527C/R527C MAD patients. For this reason, we derived MSCs from both MAD-iPSCs and healthy controls. (Note: MAD1 (age 3y) derived iMSCs patient’s data is presented or mentioned otherwise, due to the fact that youngest patient derived iPSCs retain the least epigenetic memory^13^. Though, all key experiments were reproducible in MAD2 or MAD3 derived iMSCs as well). MSCs lacked pluripotent markers, and were positive for CD105, CD90, CD73 and CD44 surface antigens (Supplementary information, Fig. S6A, B). LaminA/C expression in iMSCs increased progressively with culture duration, though at the same given passage number the MAD-derived iMSCs displayed lower protein expression (Fig. 2B-C). In addition, no pre-lamin A or progerin protein accumulation was detected in p.R527C-derived MAD-iMSCs (LMNAR527C/R527C iMSCs), as determined via western blot analysis (Supplementary information, Fig. S5A). Furthermore, the tri-lineage differentiation potential of iMSCs was conserved, but the number of lipid droplets and calcium calcification was significantly increased in MAD-iMSCs by adipogenic and osteogenic differentiation, respectively (Fig. 2D and E). The frequency of nuclear deformities, including the prevalence of donut-shaped nuclei, nuclear herniation and honeycomb structure, also increased with cell passages (Fig. 2F and G). Furthermore, increases in cell doubling time and in the expression of senescent markers, including beta-galactosidase, p16 and p21, was also associated with MAD-iMSCs (Fig. 2H-J). Taken together, these results demonstrated that iMSCs represent a suitable candidate to investigate the underlying molecular pathology of laminA/C homozygous p.R527C mutations.

### LaminA/C p.R527C iMSCs show discrete mitochondrial dysfunction, which can be rescued via correction with CRISPR/CAS9

Recently, a biallelic variant of the outer mitochondrial membrane protein metaxin-2 (MTX-2) has been found to cause mitochondrial fragmentation and result in a phenotype resembling that of patients with MAD^9^. Fragmented mitochondrial networks have also been observed in homozygous p.R527C MAD-iMSCs when stained with Mitotracker red, which was independent of excess reactive oxygen species (ROS) production (Fig. 4 A-D). It is worth noting that excess ROS production is one of the key features of HGPS-derived cell lines^1, 14^, but was not observed in MAD-iMSCs in the present study. In addition to mitochondrial fragmentation, significant reduction in ATP was also observed (Fig. 4E). This led us to examine mitochondrial membrane potential (MMP or ΔΨm), which is known to serve a vital role in regulating ATP production. JC-1 dye is a key indicator for the evaluation of ΔΨm, because in its natural state JC-1 yields green fluorescence and exists as monomer, but at higher concentrations it forms aggregates and exhibits red fluorescence^15^. The MMP of p.R527C iMSCs was found to be severely reduced when stained with JC-1 dye (Fig. 4F and G).

**Fig 4.**
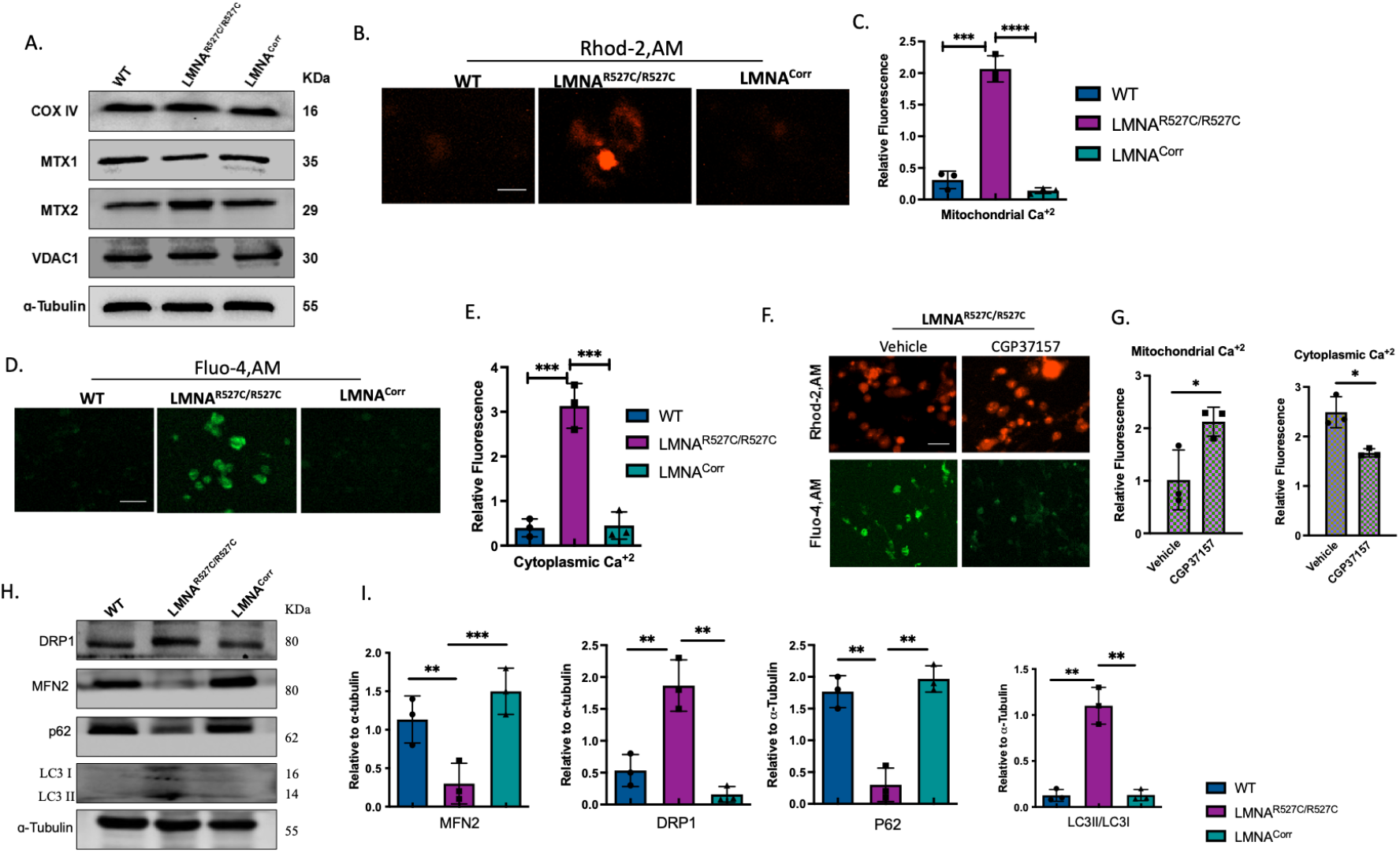
Hampered calcium homeostasis in MAD patient iMSCs. All experiments presented in this figure were in between 8 to 11cell passage number. In all cases the error bar represent standard deviation. *p<0.05, **p<0.01, and ***p<0.001, with comparisons indicated by lines. (A) Western blot analysis of whole iMSC cell lysate revealed unchanged expression of mitochondrial internal control proteins (VDAC1 and COX-IV) and mitochondrial membrane proteins (MTX1 and MTX2). (B-C) iMSCs were loaded with 0.2 µM Rhod-2,AM for 20 min in PBS, and mitochondrial Ca^+2^ levels were quantified with live imaging using a lion-x heart microscope, Ex/Em 550/585 (n=5). Scale bar represents 100 µm. (D-E) Cytoplasmic Ca^+2^ levels were measured by loading iMSCs with 2 uM Fluo-4,AM and imaging at Ex/Em 488/525 (n=4). Scale bar = 100 µm. (F-G) MAD-derived iMSCs were first incubated with 30 µM CGP37157 for 20 min followed by loading with Rhod-2,AM or with Fluo-4,AM. Decreases in the percentage of the Fluo-4 population in the CGP37157 treatment group showed that the rise in cytoplasm Ca^+2^ in MAD-iMSCs was dependent on mitochondrial calcium levels (n=3). Scale bar = 100 µm (H-I) DRP1, MFN2, p62 and LC3II and LC3I expression from whole iMSC cell lysate was analysed using western blot analysis. Bands were quantified using image J software and the expression levels were normalized to alpha-tubulin (n=4).

Although lower ATP production and decreased ΔΨm was observed in MAD-IPSC derived MSCs from all three patients, we sought to confirm the effects of p.R527C mutations effect by correcting this with CRISPR/CAS9. Mutation was biallelically corrected by delivering the RNP complex with 80 bp ssDNA donor in MAD-iPSC cells (Supplementary information, Fig. 7A-C). Homozygous corrected clones were confirmed by Sanger sequencing and then differentiated into iMSCs. The homozygous corrected mesenchymal stem cells are denoted here as (LMNA^**Corr**^). Nuclear abnormality, cell growth and senescent markers were restored in corrected iMSCs (Supplementary information, Fig. S8A-D). In addition, the inhibited ΔΨm and increased mitochondrial fragmentation was also reverted in LMNA^**Corr**^ iMSCs (Fig. 4H-K), suggesting that mitochondrial dysfunction was driven by homozygous LMNA p.R527C mutations.

### Altered Ca^+2^ homeostasis in MAD-iMSCs is the major underlying cause of mitochondrial fragmentation

Recognizing the mechanisms underlying mitochondrial dysfunction is crucial for not only characterizing the distinctive and overlapping nature of MAD with other laminopathies, but also to highlight potential targets for its rectification. To this aim, we first assessed previous studies that have investigated mitochondrial dysfunction in laminopathies. Modelling of LMNA E159K mutation in drosophila showed accelerated aging by the defective transport of mitochondrial transcripts from nucleus to cytoplasm^16^. Although we did not find differences in integral mitochondrial membrane transcripts in the nuclear fraction of MAD-iMSCs (Supplementary information, Fig. S9A-k). Levels of the outer mitochondrial membrane proteins MTX1 and MTX2 were not downregulated (Fig. 4A); this was recently reported to be the causative agent of the MAD phenotype^9^. Next, we assessed mitochondrial calcium homeostasis, which is an integral component in the maintenance of mitochondrial membrane potential dynamics^17, 18^. Cells were stained with Rhod-2AM, which emits red fluorescence directly proportional to the Ca^+2^ levels present in the mitochondria. MAD-iMSCs showed increased fluorescence compared with healthy or corrected-iMSCs (Fig. 4B-C). Increased mitochondrial Ca^+2^ levels are also known to lead to perturbed calcium levels in other cellular compartments, as the mitochondrial calcium efflux leads to increases in cytoplasmic Ca^+2^ level. Fluo-4 is a dye used to determine cytoplasmic Ca^+2^ levels. As with mitochondrial Ca^+2^ concentration, MAD-iMSCs also displayed increased cytoplasmic Ca^+2^ levels, and emitted bright fluorescence upon staining with Fluo-4 dye (Fig. 4D and E). In order to demonstrate that the rise in cytoplasmic Ca^+2^ was dependent on increased mitochondrial Ca^+2^ levels and not the other way round, we treated the cells with CGP-37157, which is a potent inhibitor of Na^+^/Ca^+2^ exchange and blocks mitochondrial calcium efflux to the cytoplasm^19^. Cells treated with CGP-37157 showed decreased cytoplasmic calcium levels (Fig. 4 F-G), confirming the dependency of cytoplasmic Ca^+2^ levels on increased mitochondrial Ca^+2^ levels in MAD-iMSCs.

Loss of mitochondrial calcium homeostasis is known to induce mitophagy, which is closely associated with the disequilibrium of mitochondrial fission and fusion^20^. Here, the induction of mitophagy was evidenced by the increased expression of LC3B and the decreased expression of p62. Along with the activation of mitophagy, expression of the mitochondrial fusion protein MFN2 was decreased, while expression of DRP1 (which is involved in peroxisomal fragmentation and mitochondrial fission) was significantly increased in MAD iMSCs (Fig. 4 H-I). Together, these results suggest that increased mitochondrial Ca^+2^ levels are the primary reason for mitochondrial dysfunction in MAD; causing the decreased ΔΨm and ATP production that eventually results in aberrant mitophagy-mediated mitochondrial fragmentation.

### Mitochondrial fragmentation in p.R527C MAD-iMSCs activates the inflammasome

As noted earlier, homozygous LMNA p.R527C patients presented with chronic inflammation which could be due to several independent factors, including DNA damage^21^, activation of transposons^22^ or the release of Mt-DNA via mitochondrial fragmentation^23^. Therefore, we aimed to systematically investigate the relationship between mitochondrial dysfunction and the inflammaging observed in patients with MAD. The expression of the proinflammatory cytokines IL-8, IL-18, IL-6 and IL-1β was increased in MAD-iMSCs (Fig. 5A), which further confirmed the involvement of MSCs in the development of inflammaging. However, the expression of IFN-δ, IFN-β and IFN-α remained unchanged (Fig. 5B), indicating that the activation of inflammatory cytokines occurred in a cGAS-stimulator of interferon genes (STING) sensor—independent manner. Interestingly, the expression of STING was significantly higher in MAD-iMSCs, but the expression of the downstream effector proteins TBK1 and p-TBK1 were remain unchanged (Fig. 5C and D). This inhibition of STING DNA sensor can be understood in the context of a recent study that linked the deactivation of STING sensor with saturated levels of calcium^24^.

**Figure 5.**
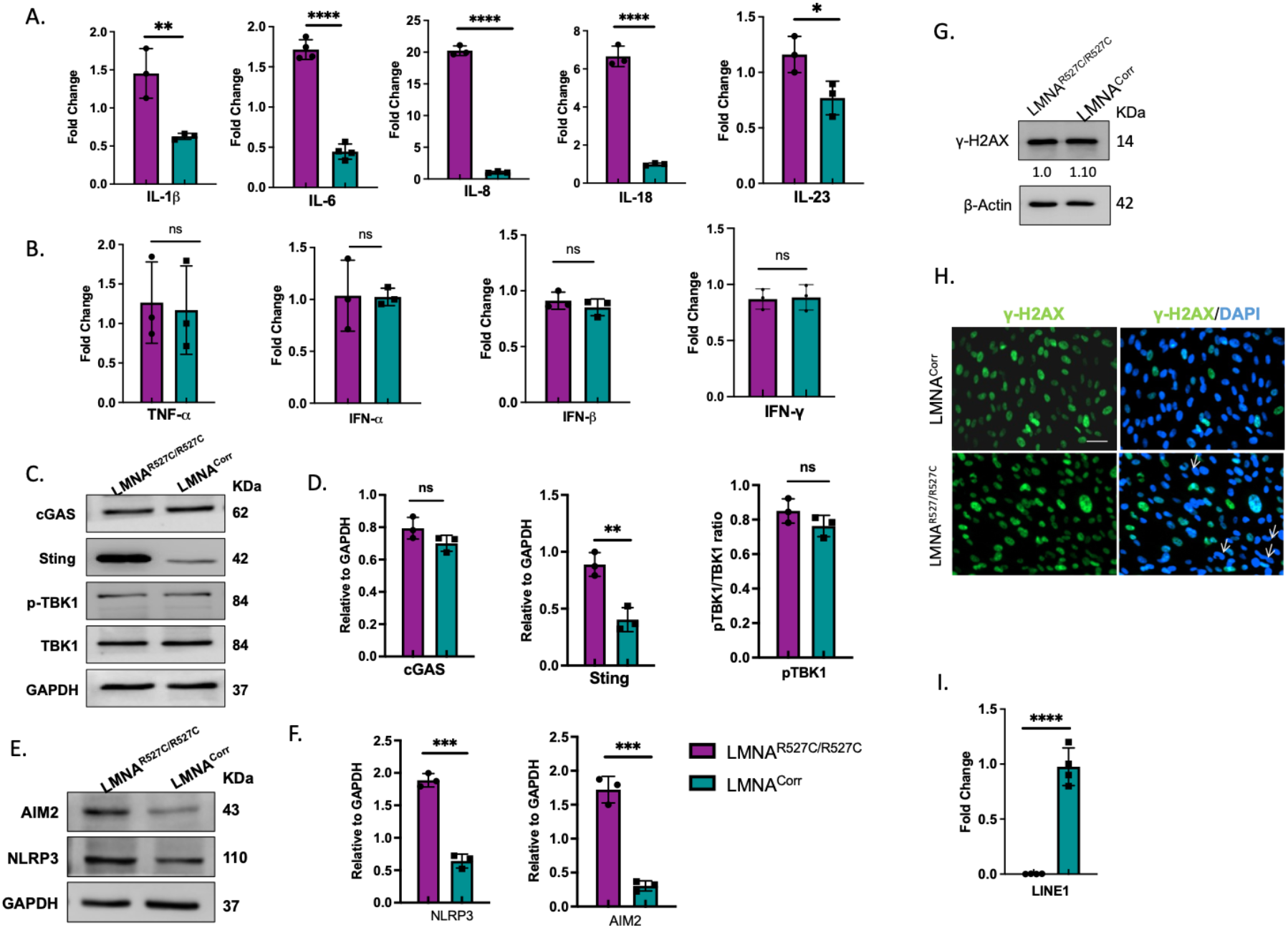
Activation of the inflammasome in MAD-iMSCs. In all cases the error bar represent standard deviation. *p<0.05, **p<0.01, and ***p<0.001, with comparisons indicated by lines. (A and B) RT-qPCR analysis of pro-inflammatory cytokines in MAD-iMSCs and corrected-iMSCs at passage 9-12 of cell culture (n=3). (C and D) cGAS, STING, TBK-1 and p-TBK1 expression in whole cell lysate was analysed by western blotting. Band intensities were quantified using image-J software and normalized with GAPDH (n=4) (E-F) AIM2 and NLRP3 expression was increased in MAD-iMSCs (n=3). (G) Representative western blot from whole cell lysate (cell passage number 11) showed the expression of γ-H2AX (n =5). (H) Immunofluorescence staining of γ-H2AX (green), nucleus (blue) was performed between 10 to 14 cell passage number (n = 4). Bar represents 200 µm. (I) RT-qPCR analysis of Line-1 from cDNA, transcribed using hexamers (n =3).

The pattern of increased pro-inflammatory cytokine expression in MAD-iMSCs indicated the activation of inflammasomes which can also be activated by other oligomeric sensors such as NLPR3 and AIM2^25^. Here, expression of absent in Melanoma 2 (AIM2) and nucleotide-binding oligomerization-like receptor family, pyrin domain-containing protein-3 (NLRP3) was increased in MAD-iMSCs (Fig. 5E and F). This could be a consequence of deregulated calcium efflux and mitophagy, which released mitochondrial DNA into the cytoplasm and so induced the inflammasome^26^. Importantly, unlike HGPS MSCs, which displayed increased expression of DNA damage markers in a previous study^11^, γ-H2AX expression did not increase in MAD-iMSCs (Fig. 5G and H). Furthermore, retrotransposons (line-1), which are known to activate DNA sensors during the natural aging process^22^, were also downregulated in MAD-iMSCs (Fig. 5I). These results suggest that mitochondrial fragmentation due to aberrant calcium regulation was the primary reason for the activation of the oligomeric sensor-mediated inflammasome, which could be responsible for the sterile chronic inflammation observed in homozygous p.R527C MAD patients.

### Activation of MAM-STAT3 deregulates Ca^+2^ homeostasis through increased IL-6 expression

After confirming that calcium-dependent mitochondrial fragmentation induces the inflammasome in MAD-iMSCs, we investigated the causative agent responsible for disrupted Ca^+2^ homeostasis. Computational analysis predicted the mutational effect of 527^th^ LMNA residue, which introduced a pocket in the outer protein structure by disrupting an arginine-glutamine salt bridge^27^, thereby disrupting the nuclear lamina and chromatin organization. Substantial intranuclear aggregation of lamin a/c, lamin B1, lamin B2 and the partial loss of emerin around the nuclear rim indicated the presence of disrupted nuclear lamin in p.R527C iMSCs (Fig. 6A). The expression of emerin, lamin B1 and B2 was also reduced, as determined via western blotting (Fig. 6B). LaminA/c is known to affect heterochromatin structure by interacting with chromatin through its topological-associated domain (TAD), as well as by activating DE acylase. To further confirm the effect of disrupted nuclear lamina on chromatin stability, we extracted chromatins and analysed histone-associated methylation and acetylation using western blotting. Several methylation events are involved in transcriptional activation, including H3K4me3 and H3K36me2/3, and these were significantly decreased in MAD-iMSCs (Fig. 6C). Similarly, altered expression of H4k20me2 and H4k20me3 was also noted (Fig. 6D). However, no difference was observed in H3K9me3 and/or H3K27me3/H3K27ac3 levels (Fig. 6E), although altered methylation and acetylation in H3k9/H3k27 is a characteristic feature of HGPS-derived fibroblasts and MSCs^28^. This suggested that epigenetic landmarks of HGPS (representing typical progeria) and MAD (atypical progeria) did not result in similar heterochromatin modification. Moreover, expression of nuclear deacylases (sirt1, sirt6 and sirt7), which directly interact with lamin a/c, was also downregulated in MAD-iMSCs (Fig. 6F).

**Figure 6.**
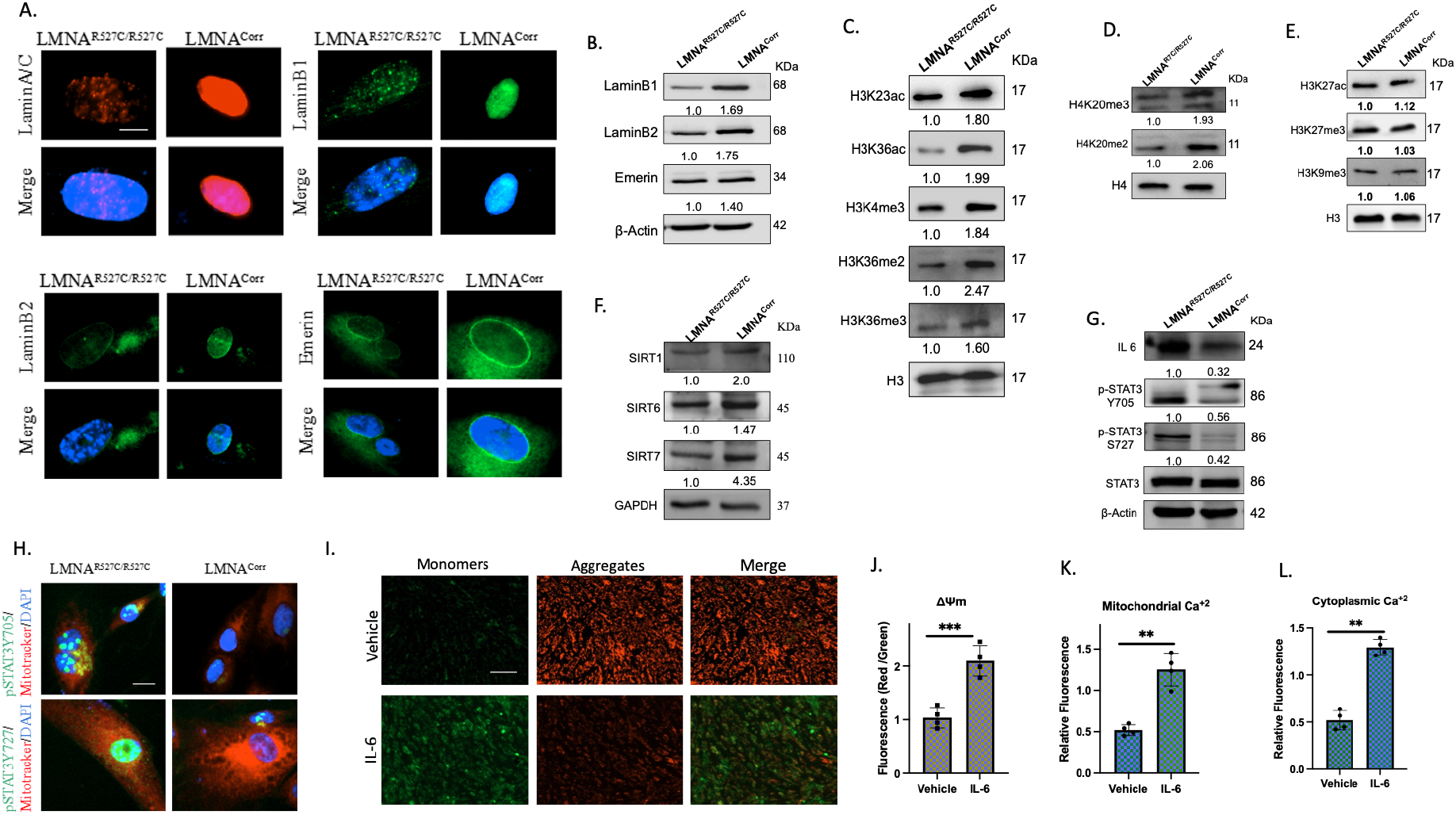
Disrupted nuclear lamina and heterochromatin remodeling activates MAM-STAT3 in MAD MSCs. In all cases the error bar represent standard deviation. *p<0.05, **p<0.01, and ***p<0.001, with comparisons indicated by lines. Cell passage number = ≤ 10. (A) Representative images show the immunofluorescent staining of LMNA (red), LaminB1 (Green), LaminB2 and Emerin (Green), with nuclei stained with DAPI (Blue). Scale bar = 20 µm. (B) Western blot analysis from whole cell lysate showed the decreased expression of the nuclear lamina proteins laminB1, LaminB2 and Emerin in MAD-iMSCs. (C-E) Western blot analysis was performed after chromatin extraction from MAD-iMSCs and corrected-iMSCs. H3 and H4 represents the internal controls (n=3). (F) Western blot analysis of whole cell lysate showed decreased expression of nuclear sirtuin proteins, Sirt1, Sirt6 and Sirt7 (n=4). (G) Western blot analysis from whole cell lysate revealed the increased expression of iL-6, and activated p-STAT3^y705^ p-STAT3^s727^ (n=3). (H) Immunofluorescent staining showed the co-localization of p-STAT3^y705^ with the mitochondrial associated membrane, while p-STAT3^s727^ was found to be localized in the nucleus in MAD-iMSCs. Scale bar = 20 µm. (I and J) Wild type iMSCs were treated with 20 ng/ml IL-6 for 24 h and evaluated for mitochondrial membrane potential. Scale bar = 200 µm. (K and L) Quantification of mitochondrial and cytoplasmic calcium levels after treating cells with 20 ng/ml IL-6 using Rhod-2,Am and Fluo-4,Am respectively.

**Figure 7.**
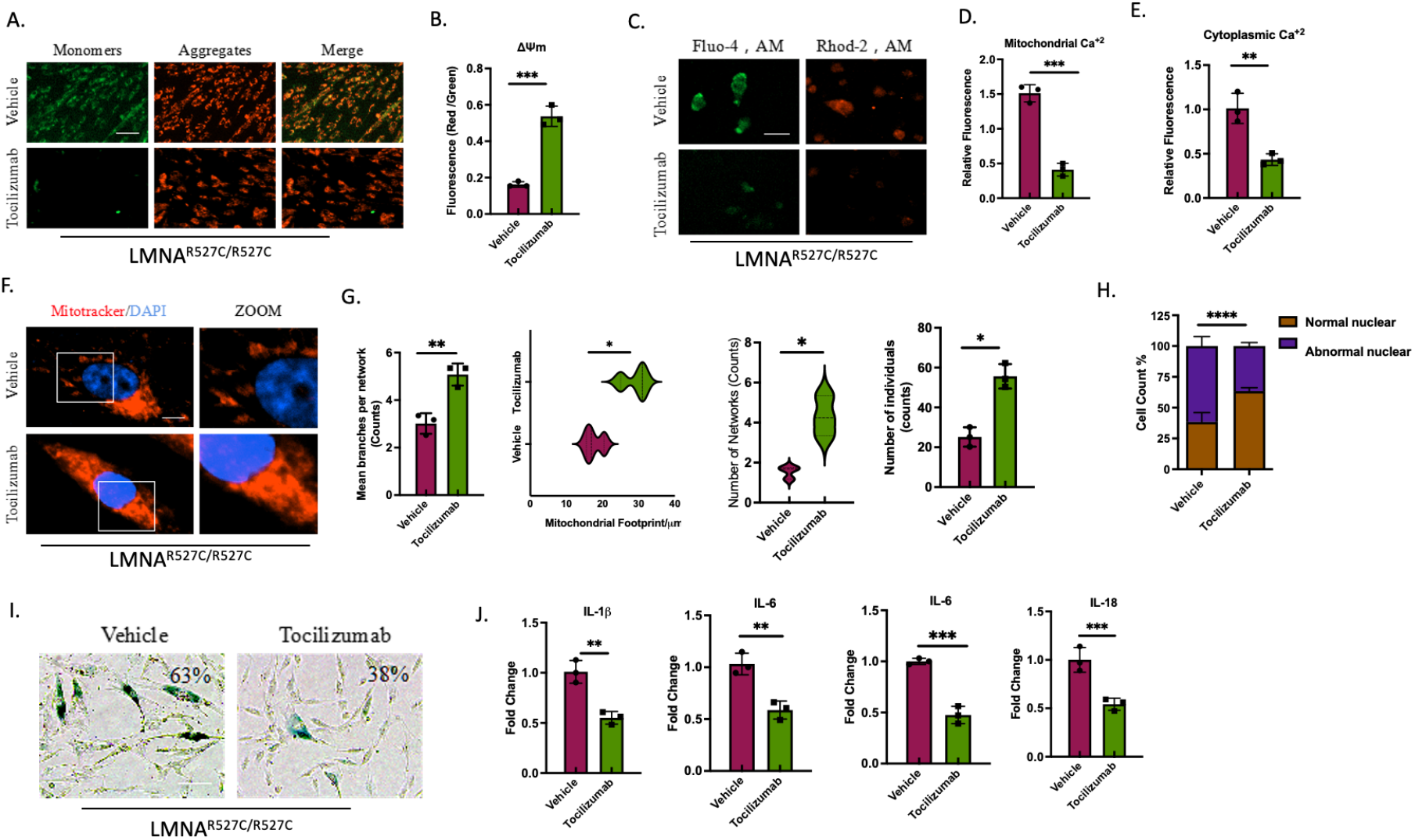
STAT3 inhibitor rescues nuclear and mitochondrial defects associated with LMNA p.R527C mutations. All experiments presented in this figure were in between 9 to 13 cell passage number. In all cases the error bar represent standard deviation. *p<0.05, **p<0.01, and ***p<0.001, with comparisons indicated by lines. (A and B) MAD-iMSCs were treated with 10 µg/ml Tocilizumab for 24 h period. JC-1 staining revealed that cells treated with Tocilizumab regained their losst mitochondrial membrane potential. Scale bar = 200 µm. (C-E) Cells treated with tocilizumab for 24 h followed by quantification of Mitochondrial and cytoplasmic Ca^+2^ levels with Rhod-2,AM and Fluo-4,AM dyes at Ex/Em 4550/585 and 488/425 nm, respectively. (F and G) Mitotracker staining on Tocilizumab-treated cells (post 7 days) showed decreases in mitochondrial fragmentation with increases in mitochondrial mean branch per network. (G) Beta-galactosidase staining was performed after 21 days of tocilizumab (10 µg/ml) treatment. Scale bar = 200 µm (n=8). (H and I) MAD-iMSC nuclei were stained with DAPI after 21 days of with Tocilizumab treatment. Scale bar = 20 µm (n =8). (J) RT-qPCR analysis of pro-inflammatory cytokines in MAD-iMSCs treated with tocilizumab for 48 h.

These alterations to heterochromatin should be responsible for a substantial degree of the differential gene expression observed in MAD, though it would be wise to look for other candidates that might be involved in the deregulation of calcium homeostasis. For example, downregulation of nuclear sirtuin, including sirt1 and sirt6, is also known to increase the expression of IL-6^29, 30^. The upregulation of IL-6 in serum taken from MAD patients (Fig. 1D), as well as in MAD-iMSCs, drew our attention to a recently-defined, noncanonical role of stat3. Upon activation, stat3 has been shown to relocate on the mitochondrial associated endoplasmic reticulum membrane (MAM) and modulate Ca^+2^ homeostasis^31, 32^. In the present study, there was a significant increase in both Tyr^705^ and Ser^727^ phosphorylation in patient MAD-iMSCs (Fig. 6G), which denoted the activation of the JAK/Stat pathway. Notably, only Tyr^705^ Stat3 was found to localize to the MAM when co-stained with Mitotracker, while p-Ser^727^ Stat3 showed higher but more uniform nuclear expression (Fig. 6H). To further confirm the role of MAMs-stat3 in destabilization of Ca^+2^ homeostasis, stat3 was activated by exogenous IL-6 treatment in healthy iMSCs, and then tested for the change in MMP and levels of mitochondrial and cytoplasmic Ca^+2^. Deterioration of MMP, along with increases in mitochondrial and cytoplasmic Ca^+2^, was observed in normal iMSCs treated with 20 ng/ml IL-6 (Fig. 6I-L). These results suggested that MAM-stat3 was involved in the regulation of Ca^+2^ homeostasis, and contributed to mitochondrial dysfunction.

### Inhibition of MAM-Stat3 rescues Ca^+2^ homeostasis, the expression of pro-inflammatory cytokines and senescence in MAD-IMSCs

Since Stat3-Tyr^705^ was found to be localized on the MAM, we next investigated whether inactivating MAM-stat3 could rescue mitochondrial dysfunction and early senescence in MAD-derived iMSCs. To this end, we used four different FDA-approved Stat3 blockers, which dephosphorylate Stat3 in different ways. Cells were treated with various concentration of Tocilizumab (an anti-human IL6-R/CD-126 antibody), LMT-28 (a synthetic inhibitor for IL6-R-Beta/gp130/CD130), tofacitinib (a JAK family inhibitor) and AZD8055 (an m-TOR inhibitor), and assessed for mitochondrial Ca^+2^ levels and mitochondrial membrane potential. Ca^+2^ homeostasis and ΔΨm was best rescued by Tocilizumab (Fig. 7A-E). Other STAT3 inhibitors had also rescued mitochondrial dysfunction (supplementary information, Fig. S10A-D). The change in ΔΨm and Ca^+2^ homeostasis induced by Tocilizumab rescued not only decreased β-galactosidase staining intensity, but also decreased nuclear dysmorphism and mitochondrial fragmentation in p.R527C iMSCs (Fig. 7F-I). Moreover, the mRNA levels of other pro-inflammatory cytokines including IL-8, IL-1β and IL-18 were also reduced and mitochondrial function was improved, suggesting that the inflammasome was inactivated (Fig. 7J). Overall, these results strongly suggest that mitochondrial integrity serves an intermediary role in the development of progeroid syndrome, and inactivation of MAM-stat3 by FDA-approved inhibitors would be a reasonable option for treating p.R527C MAD patients.

### MAD-iMSC extracellular vesicles lose their immunomodulatory capabilities and aggravate inflammaging at a systemic level

MSCs regulate the homeostasis of neighbouring tissues, distant organs and immune cells by various cytokines, chemokines and growth factors that are secreted as encapsulated extracellular vesicles (EVs). Emerging evidence suggests that age-related changes in MSCs alter the composition of EVs, which in turn accelerates aging at the systemic level^33^. Thus, we investigated the restorative effect of MAD-iMSC EVs on the well-developed rodent bleomycin-induced lung fibrosis model. Since patients affected with systemic sclerosis (SSC) often present with internal organ failure and fibrosis^34^, the SSC and chronic inflammation observed in patients with MAD (Fig 1C to G and supplementary information, Fig S2) suggests that these internal organs may also be affected by fibrosis.

EVs were purified from iMSC serum-free culture medium using commercial columns. Purified EVs were enriched with EV marker, and an average diameter of 180 nm was recorded with a Nanosight particle analyser (supplementary information Fig. S11). Only EVs from healthy control iMSCs rescued the bleomycin-induced aberrant extracellular matrix deposition in the mouse lungs, as determined via trichome or picrosirius red staining. On the contrary, MAD-iMSC EVs yet enhanced collagen deposition and worsened the fibrotic score compared with the vehicle control (Fig. 8A-D). This aggravating effect of the MAD-iMSC secretome was further evaluated by utilizing an in vitro hydrogel 2D cell culture system. Hydrogel biomaterials are now increasingly utilized in the in vitro culture system by governing substrate stiffness, thereby recapitulating the physiological microenvironment^35^. Several studies have shown that fibroblast cells, cultured on lower stiffness or elastic modulus <5kPa remain quiescent, but differentiate into a diseased phenotype on stiffer substrate (E>20 kPa)^36^. Here, we cultured fibroblasts on 3% gelatin methacryloyl (GelMa) hydrogel, which represents normal lungs stiffness (4 kPa)^37^ and were further treated with p.R527C iMSCs and control EVs. Fibroblasts treated with MAD-iMSCs showed increased expression of fibrogenic myofibroblast markers and significant increases in soluble collagen content, indicating the transformation of fibroblasts towards fibrosis (Fig. 8E and F). Interestingly, increased expression of IL-6 was also detected in EVs derived from MAD-iMSCs, and this has the ability to alter the ΔΨm of healthy cells (Fig. 8G and H). From these results we can infer that MSCs, and similar cells such as fibroblasts with p.R527C mutation, might also be participating in mitochondrial damage-mediated senescence and inflammation. Accordingly, the proposed IL-6 inhibitors would be beneficial in restoring the systemic inflammation instigated by cells of mesenchymal origin in patients with MAD.

**Figure 8.**
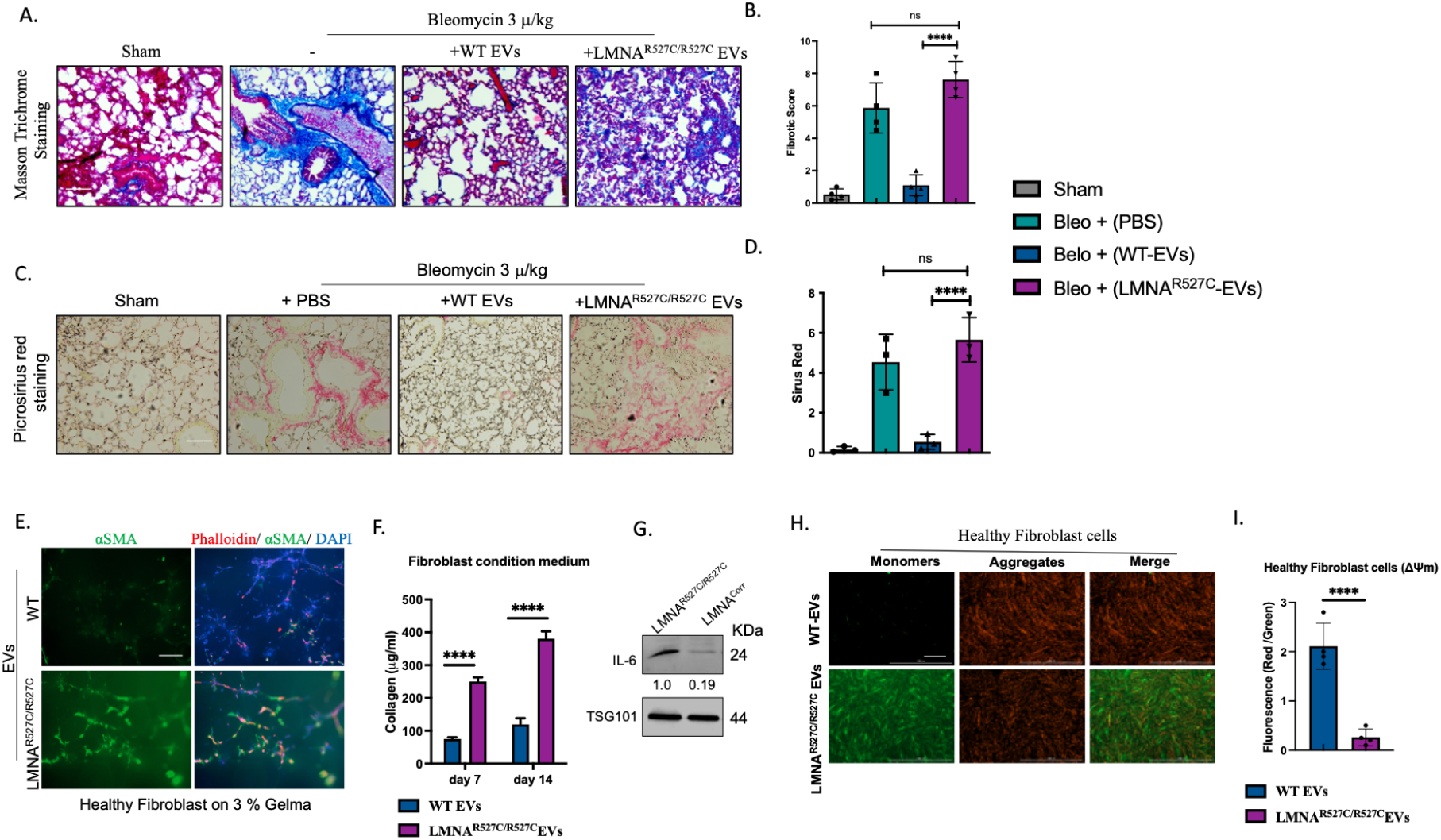
Fibrosis-provoking effects of extracellular vesicles from MAD-iMSC. In all cases the error bar represent standard deviation. *p<0.05, **p<0.01, and ***p<0.001, with comparisons indicated by lines. (A) Paraffin (5 µm)-embedded lung tissue was sliced and stained with trichome-masson dye. Scale bar = 100 µm. (B) Quantification of Masson trichome staining. Fibrotic score was calculated on the basis of collagen content deposition and architectural destruction by two professional histologists that were blinded to the treatment. LMNA^R527/R527C^ exosomes could not rescue bleomycin induced fibrosis but enhanced the fibrotic score compared to group treated with PBS only, though it was non-significant. (C and D) Sectioned lung tissues were stained with Picrosirius dye, which showed collagen fibres in red. Scale bar = 100 µm. (E) Normal fibroblast cells were cultured on 3% Gelma hydrogel (as described in the methods section) and treated with MAD or WT MSC-derived EVs (1×10^6^) every fourth day. IFC was performed on the 14^th^ day of cell seeding. Increased alpha sma expression indicated the induction of fibrosis. Scale bar 200 µm. (F) Soluble collagen was quantified in fibroblast culture medium supernatant using sircol assays. (G) Western-blot of EVs derived from patient iMSCs or healthy control. (H and I) Healthy fibroblast cells were treated with MAD-iMSC exosomes for 24 h and then evaluated for mitochondrial membrane potential using JC-1 dye. Scale bar = 200 µm.

## Discussion

Age-associated diseases share some basic pathological features, typically concerning inflammation. In the present study, we observed immunosenescence in patients with homozygous LMNA R527C mutations, and systematically characterized the underlying molecular pathology responsible for this immunosenescence. Analysis of patient IPSC-derived iMSCs revealed that increased mitochondrial Ca^+2^ levels were the primary cause of mitochondrial dysfunction and subsequent fragmentation, which is followed by inflammasome formation. Our results suggest that patient iPSC-derived mesenchymal cells may represent a more viable model for studying the underlying pathology of such laminopathies compared with rodent models. For MAD, transgenic mice with LMNA R527C and R527H mutations do not significantly differ from the wild type in terms of lifespan. Previously, our collaborators demonstrated that DNA damage mediated premature aging in R527C transgenic mice only when these were subjected to a high-fat diet (HFD)^38^, although the DNA damage maker γ-H2AX in patient-derived R527C cells remained unchanged. In contrast, a HGPS LMNA^G609/G609^ mouse model showed twice the lifespan when subjected to an HFD compared to the group that fed with regular chow^14^. Therefore, extra care is needed in the selection of laminopathy-associated rodent models. Since the mesenchymal stem cells derived from IPSCs showed major defects associated with HGPS^11, 12^, this model maybe more useful when attempting to investigate laminopathies, such as those seen in patients with homozygous LMNA p.R527C mutations.

Severe bone abnormalities, including localized osteoporosis and generalized osteolysis, are commonly observed in MAD. The balance of osteogenic and lipogenic MSC differentiation controls bone remodelling^39^. In this study, we observed aggressive adipogenesis and premature osteogenesis in MAD-derived iMSCs, which is in line with the results of a previous study, in which ectopically-expressed R527C LMNA lead to increased osteogenesis via NOTCH signalling^40^. This premature osteogenesis could be responsible for abnormal bone density seen in patients with MAD, although it is also likely that the acro-osteolysis observed in such cases is largely due to the excess production of osteoclasts. IL-9 expression, which is persistently high in patients with R527C-associated MAD (Fig. 1D), has been shown to promote osteoclast formation in rheumatoid arthritis^41^. Furthermore, IL-6 has been shown to activate osteoclastogenesis by regulating RANKL via the JAK-STAT3 pathway^42^. For this reason, we suggest that IL-6 inhibitors may represent a viable therapeutic option to decrease bone resorption in patients with MAD. In the future, IPSC-derived osteoclasts may be of great benefit during drug screening, and may help aid our understanding the acro-osteolysis and lipodystrophy that occurs in MAD.

Disrupted nuclear lamina was observed in R527C derived cells without the accumulation of pre-lamin-A protein, despite the accumulation of progerin or pre-lamin-A being considered a common attribute of HGPS and MAD disorders, respectively ^43^. Thus, we can infer that heterochromatin modification caused by disrupted nuclear lamina could occur independently from pre-lamina-A. Heterochromatin modification caused by disrupted nuclear lamina in MAD shares some common patterns with HGPS, but some features are unique. For example, decreased histone markers for H3k9me3 or H3K27me3 is observed in HGPS, but remains unchanged in p.R527C-MADA. Nuclear sirtuin knockout mice (including sirt1, sirt6 and/or sirt7 alone) have also been shown to develop progeroid features^44, 45, 46^. The decreased expression of nuclear sirtuin observed in iMSCs in the present study could be due to the loss of direct interactions with mutant laminA/C, or indirectly. For instance, Set7-N-lysine Methyl transferase inversely regulates the expression of sirt1, and was found to overexpressed in MADA-iMSCs (Supplementary information fig.s12). In addition, chromatin remodelling in laminopathies also activates common downstream transduction pathways that are associated with mitochondrial dysfunction.

Emerging evidence suggests that dysfunctional mitochondrial metabolites relay signals to the nucleus via mito-nuclear communication, which further alters the epigenetic regulation of genes associated with aging^47, 48^. Thus, the observed alterations in epigenetic modifications in HGPS and MADA may not solely be due to disruptions in the nuclear lamina, and may also be impacted indirectly by mitochondrial dysfunction. Treating HGPS cells with the mitochondrial-targeting antioxidant methylene blue not only rescued mitochondrial defects, but also rescued nuclear abnormalities^49^. Furthermore, primary defects in mitochondria with MTX2 mutations also include nuclear blebbing^9^, reflecting the impact of mitochondrial stress on the integrity of the nuclear membrane. In the future, using techniques such as ATAC-seq to study epigenetic landmarks in HGPS or MADA-derived cells treated with drugs targeting the mitochondria would likely elucidate the impact of lamin a/c mutation and/or dysfunctional mitochondria on cellular function.

In the present study, we have systematically observed different mitochondrial stressors that have been observed in various laminopathies, and demonstrated that IL-6-dependent deregulation of intracellular Ca^2+^ levels represents a novel root cause of mitochondrial dysfunction. Previously, collagen-induced arthritis mouse models were found to display severe ankle joint stiffness, which was caused by aberrant Ca^+2^ signalling and IL-6-mediated NLRP3 inflammasome formation^50^. Ulcerated and stiff ankle joints was observed in the patients described here, as well as in previously reported patients with LMNA p.R527C MAD^10^. This suggests that aberrant Ca^+2^ signaling may be the chief cause of this phenotype. Interestingly, in serum from both MAD patients (with different LMNA mutations) and HGPS patients, IL-6 levels remain unchanged^51, 52^. However, increased intracellular Ca^+2^ levels and the differential expression of calcium regulators, including ITPR, CACNA2D1 and CAMK2N1, have been found in HGPS cells^53^. Similarly, the upregulation of CACNA1A, which regulates P/Q-type calcium channel, has been shown in haploinsufficient mutation models for lamin a/c^54^. Therefore, it is probable that STAT-3 may not be the only factor underlying deregulated Ca^+2^ signalling, and the aforementioned Ca^+2^ regulators may have additive or synergistic effects during the destabilization of mitochondrial Ca^+2^ homeostasis. Examination of these Ca^+2^ channels in the context of laminopathies may represent a promising direction for future study.

The mitochondria orchestrate a wide range of cellular processes, including energy production, cell metabolism, cell death and inflammation. The release of mitochondrial DNA (mt-DNA) into cytoplasm or the extracellular milieu is sensed as “foreign” DNA, and activates different innate immune response including TLR9, cGAS-STING and inflammasome formation^23^. Inflammasomes are multimeric protein complexes which activate the IL-1B and IL-18 mediated inflammatory form of cell death, called pyroptosis^55^. Previous studies have demonstrated that Ca^+2^ mobilization acts as a potent trigger of mitochondrial dysfunction and NLRP3 inflammasome formation via mitochondrial-associated ligands^56, 57^. This NLRP3 activation contributes to a vascular inflammatory response that leads to the development of atherosclerosis^58^, which was observed in patients with bearing homozygous p.R527C mutation.

Multiple cellular processes and organs are affected in laminopathies and, as with other diseases that involve the dysfunction of multiple systems, it is likely that a combination of therapies will be needed to treat MAD successfully. The direct correction of mutations by CRISPR/CAS9 has yielded promising results in a mouse model^59, 60^, but there remains a long way to go before this can be applied to humans. Utility of stem cell-derived EVs are currently active in clinical trials for various aging disorders. We have shown that MAD patient derived EVs has lost the immunomodulatory effect. On that account, injection of EVs derived from IPSCs or MSCs carried immense potential for increasing the lifespan of HGPS and/or MAD patients. Due to patient scarcity, <6% of all rare diseases have approved treatment options, and may be overlooked due to lack of appropriate models. However, these patients still require treatment. Thus, the repurposement of clinically-approved IL-6 or STAT-3 inhibitors, for which dosage is well defined, may represent a promising treatment for patients with homozygous p.R527C-associated MAD.

## Methods

### Consent

The experiment conducted in the study was approved by Shenzhen Medical Ethics Committee, and Shenzhen Luohu People’s Hospital Medical Ethics Committee, China. Written informed consent for the patients obtained from their parents, signed by themselves.

### Genotyping

Genomic DNA was isolated from PBMCs using DNeasy Blood & Tissue kits, according to the manufacturer’s protocol (Qiagen, Hilden, Germany). Direct sequencing of common genes involved in progeroid syndromes, including both introns and exons of LMNA, ZMPSTE24, POLD1 and BNF1, was conducted in probands, siblings and parents.

### Flow cytometry

Cytokine levels in the plasma and cell culture medium were detected using the LEGENDplex™ Human Th Panel (BioLegend, San Diego, CA, USA) containing 13 human antibodies targeting IL-5, IL-13, IL-2, IL-6, IL-9, IL-10, IFN-γ, TNF-α, IL-17A, IL-17F, IL-4, IL-21 and IL-22. Full antibody details are provided in the “Antibodies” section. Samples from healthy controls were used for cross-comparison. Cells were harvested and blocked with 1% fetal bovine serum (FBS,cat.no.10100147,Thermo Fisher scientific) for 20 min at room temperature (RT). Cells were then labelled with fluorescein-conjugated antibodies (see “Antibodies” section for details) for 30 min at RT in 1% FBS. Following incubation, cells were washed three times with 1% FBS. Flow cytometry was performed using a Flow Cytometer (Beckman Coulter Inc., Brea, CA, USA) and analysed with FlowJo software (BD Biosciences, Franklin Lakes, NJ, USA).

### Isolation and culture of PMBCs

Whole blood was collected in heparin-coated tubes and centrifuged using Ficoll gradient for 45 min at 900 x *g*. PBMCs were washed with PBS prior to suspension in PBMC culture medium, which comprised of StemSpan SFEM II medium with Stemspan CD34^+^ supplement (STEMCELL Technologies, Inc., Vancouver, Canada). Cells were maintained for 4 days at 5% CO^2^ at 37°C before further use.

### Generation and characterization of IPSCs

Cultured PBMCs were counted, and 3×10^5^ cells were transferred to 1 ml fresh complete PBMC medium and transduced with Yamanaka factors using a non-integrated CytoTune®-iPSC2.0 Sendai Reprogramming kit, (Life Technologies; Thermo Fisher Scientific, Inc., Waltham, MA, USA; cat. No. A16517). Klf4, Oct4, and Sox2 (KOS) were packaged in the same vector and transduced at a multiplicity of infection (MOI) of 5; simultaneously c-Myc and the Klf4 virus which were packaged separately was transduced at an MOI of 5 and 3, respectively. After transduction for 24 h, cells were seeded in 12-well plate coated with vitronectin (STEMCELL Technologies, Inc.). Stemspan medium was replaced with MTesR complete medium following 72 h transduction, and changed every other day for 21 days. IPSC-like colonies were manually selected by marking the outline with a 28 cc gauge syringe needle, and were then transferred to new vitronectin-coated plate using a 200 µl pipette. Different colonies from each donor were further characterized using Giemsa staining, alkaline phosphatase staining, or by teratoma formation in nude mice. In brief, cells were split in 1:10 to 1:50 ratios using ReLeSR, which is a gentle dissociation reagent, or with Accutase (both obtained from STEMCELL Technologies, Inc.). Then, cells were resuspended in mtesR-plus medium with 10 µm Y-27632 (STEMCELL Technologies, Inc.) for 24 h before this was replaced with complete MTesR plus medium.

### Karyotyping

Karyotype analysis was performed in the cytogenetic lab of Shenzhen University (China). Chromosome resolution additive and 10 mg/ml colchicine were added to the cells for 60 min at 37 °C. Cells were then harvested and treated with 0.075 M KCl solution for 30 min. Slides were fixed with a 1:3 ratio of acetic acid:methanol and analysed for Giemsa banding.

### MSC differentiation

Human iPSC clones were differentiated into mesenchymal stem cells (MSCs) using a Stemdiff Mesenchymal Progenitor kit, according to the manufacturer’s protocol (STEMCELL Technologies, Inc.). Briefly, iPSCs at 50% confluence were induced with early mesodermal progenitor medium (STEMdiff-ACF) for 4 days, followed by two days incubation with MesenCult-ACF Medium. Cells were passaged at day 6 on MesenCult-ACF Attachment Substrate-coated wells in MesenCult-ACF medium. Cells were split once they reached 80% confluency and characterized 20-22 days after differentiation by qPCR, flow cytometry and the standard tri-lineage differentiation.

### Tri-lineage differentiation

The tri-lineage potential of iMSCs (into adipocytes, osteocytes and chondrocytes) was evaluated with MSCgo-adipogenic, MSCgo-osteogenic and MScgo-chondrogenic differentiation kits (all from Biological Industries Ltd., Cromwell, CN, USA) according to the manufacturer’s protocol. Briefly, 6×10^4^ cells/well were seeded in 0.1% gelatin-coated 24-well plates and treated with complete adipogenic medium or osteogenic medium for the indicated time, before being fixed with paraformaldehyde. In addition, 2×10^5^ cells/well were seeded in U-bottom 96-well plates, and were used to differentiate iMSCs in chondrocytes.

Early osteogenic potential was evaluated with alkaline phosphatase (Beyotime Biotechnology, China) on day 7, while calcium nodules were stained with Alizarin red staining dye (Vivacell Biotechnology GmbH, Denzlingen, Germany; cat. no. MC370-1.4) after 21 days of osteogenic differentiation. Oil Red-O (Vivacell Biotechnology GmbH; cat. no. MC37A0-1.4) and Alcian blue staining (Vivacell Biotechnology GmbH; cat. no. MC37B0-1.4) was also performed to assess the formation of lipid oil droplets and cartilage, respectively.

### Senescence Associated Beta-Galactosidase (SA-β-Gal) staining

SA-β-Gal staining was performed according to the manufacturer’s protocol (Beyotime Biotechnology, China). Briefly, cells were seeded in six-well plates at a density of 30-40%, and were fixed at the indicated passage number or treatment once they reached 70-80% confluency. After washing with PBS, cells were stained overnight with freshly prepared SA-β-Gal staining solution at 37°C. Images were acquired with brightfield using a Lionheart FX automated microscope (BioTek; Agilent Technologies, Santa Clara, CA, USA).

### Western blot analysis

Cells were lysed in RIPA lysis buffer (Beyotime Institute of Biotechnology) containing protease and phosphate inhibitors (MedChemExpress LLC, Monmouth Junction, NJ, USA). Protein concentration was determined using a BCA kit (Thermo Fisher Scientific, Inc.). Lysates were denatured with 6X SDS loading buffer for 10 min at 98°C, and 20 µg total protein was run on each lane of an SDS PAGE gel. After the protein was transferred to a PVDF membrane (Millipore; Merck KGaA, Darmstadt, Germany; cat. no. IPVH00010), target proteins were detected using the antibodies mentioned in the “Antibodies” section and (Supplementary information Tables S9-S1). Images were captured using a ChemiDoc Touch system (Bio-Rad Laboratories, Hercules, CA, USA) and band intensities were quantified using Image J software (version 1.8.0; National Institutes of Health, Bethesda, MD, USA).

### Immunofluorescence

IPSCs and MSCs, seeded 48 or 24-well plates and that had reached the confluency between 50 to 80%, were fixed with 4% paraformaldehyde at RT for 15 min, permeabilized with 0.1% Triton X-100, and then blocked with 5% BSA for 1 h at RT. Cells were incubated overnight with primary antibodies at 4°C. After washing with PBS, cells were stained with fluorescently-labelled secondary antibodies for 45 min at RT (see “Antibodies” section for further details). Cells were then stained with phalloidin (1:1000 dilution; Abcam, Cambridge, UK) and/or 1 µg/ml DAPI (Sigma-Aldrich; Merck KGaA) for 5 min at RT. Images were acquired using a Lionheart FX Automated Microscope (BioTek; Agilent Technologies).

### EV purification and characterization

Cell conditioned medium was collected at 80% cell confluency. Debris and floating cells were removed by centrifuging at 1000 x *g* for 15 min, followed by passing the medium through a 0.22 µm filter (Millipore; Merck KGaA). Cell supernatant (50 mL) was then concentrated to 5 mL by centrifugation using Amicon® Ultra-15 Ultrafiltration Centrifuge Tubes (Millipore; Merck KGaA) following the manufacturer’s protocol. Isolation of exosomes was carried with concentrated supernatant using qEV size exclusion columns/70nm (Izon Science Ltd., Christchurch, New Zealand), according to the manufacturer’s instructions. Purified exosomes were aliquoted and immediately stored at - 80°C. The size distribution of isolated EVs was determined using a NanoSight Tracking Analysis NS300 system (Malvern Panalytical, Malvern, UK). The Zeta potential and polydispersity index of EVs were obtained using a Zetasizer Nano ZS molecular size analyzer (Malvern Panalytical).

### *In vivo* bleomycin-induced lung model

All animal experimental procedures and maintenance abided by the National Research Council’s Guide for the Care and Use of Laboratory Animals. All procedures were approved by the Institutional Animal Care and Use Committee of Shenzhen University (Shenzhen, China). ∼30 Male C57BL/6 strain mice (10-8 weeks old; 20-22 g) were purchased from Vital River Laboratory Animal Technology Co. Ltd, (Beijing, China) and were housed at 22 ± 5 °C in a 12 h light/dark cycle and supplied with food and water *ad libitum*. Mice were randomly divided into bleomycin-treated and sham groups. Anaesthesia was maintained by inhalation of 4% isoflurane while a single endotracheal dose of bleomycin sulphate (50 μl, 3 U/kg) or 50 µl of PBS as a sham control was administered to each individual. Furthermore, mice treated with bleomycin received a single tail vein injection at day 0 and at day 7 of either 200 µl PBS, EVs derived from MAD-iMSCs, or EVs derived from WT-iMSCs. Each EV dose contained ∼1 × 10^9^ particles per 200 µl volume. Each treatment group have six mice in total. The mice were euthanized 14 days after the initial administration of bleomycin or the sham treatment, and underwent histological examination.

### Synthesis and preparation of GelMA hydrogels

GelMA hydrogels were synthesized according to a previously reported protocol^61^. Briefly, 20 g gelatinA (Sigma-Aldrich; Merck KGaA) was added into 100 ml of phosphate buffer, and the solution was maintained at 60°C with constant stirring until the gelatinA was completely dissolved. Next, 2 ml methacrylic anhydride (Sigma-Aldrich; Merck KGaA) was added dropwise with vigorous stirring, and the mixture was maintained at 60°C for a further 3 h. The reaction mixture was then dialyzed using a 12-14 kDa membrane (Solarbio Life Science, Beijing, China), and subsequently freeze dried.

GelMA hydrogels with different degrees of stiffness were obtained by adding the 2.5, 5, 10, and 15% (w/v) GelMA powder to Dulbecco’s PBS (Gibco; Thermo Fisher Scientific, Inc.) with 0.5% (w/v) of a water-soluble photo-initiator (Irgacure 2959; Sigma-Aldrich; Merck KGaA). The solution was then mixed and filtered through a 0.22 μm filter. Next, ∼150 µl of the solution was added to 48-well culture plates to cover the surface area, and the plates were then exposed to UV light (360 nm; Run LED, China) for 30 secs. GelMA hydrogels were washed twice with PBS prior to seeding with cells.

### Mitotracker red staining and mitochondrial network and fragmentation analysis

Cells were seeded at a density of 3×10^4^ cells per well in 48-well plates, and incubated with 20 nM of mitotracker red (ThermoFisher Scientific, Inc.) for 30min at 37°C. Cells were washed twice and then fixed with paraformaldehyde, following which nuclei and actin were stained with DAPI and phalloidin, respectively, at 1 µg/ml for 5 min at RT. The resultant images were processed as described earlier^62^, using the freely available Mitochondrial network analysis (Mina) toolset (https://github.com/ScienceToolkit/MiNA,it.)

### ROS production

iMSCs were seeded at a density of 10,000 cells per well in 48-well plates for 24 h, and incubated with DCFH-DA Reagent (Beyotime Institute of Biotechnology) at final concentration of 10 µM for 30 mins at 37°C. The cells that served as a positive control were incubated for 25 min at 37°C with Rosup at a dilution of 1:1000 prior to staining with fluorogenic DCFH-DA dye. Images were acquired at Excitation/Emission (Ex/Em) 488/525 nm using a Lionheart FX Automated Microscope (BioTek; Agilent Technologies).

### ATP quantification

ATP was quantified on lysed cells using a luminescent ATP detection kit (Abcam), following the manufacturer’s protocol. Bioluminescence was measured using a Biotek H1 plate reader (BioTek; Agilent Technologies).

### Mitochondrial membrane potential (MMP) assay

The MMP (ΔΨm) of cells was evaluated using the cationic fluorescent dye JC-1 (Abcam). Briefly, cells cultured in either 24 or 48 well plates were incubated with JC-1 dye for 20 min at 37°C, and observed through green and red channels with Ex/Em of 488/525 and 550/585 nm, respectively, using a Lionheart FX automated microscope (BioTek; Agilent Technologies). Fluorescence intensity was further quantified using Image J software (ImageJ 1.8.0; National Institutes of Health, Bethesda, MD, USA).

### RT-qPCR

Nuclear and cytoplasmic fractionation was conducted using the Nuclear and Cytoplasmic Extraction Reagents kit (Beyotime Institute of Biotechnology) according to the manufacturer’s protocol. Total RNA, either from whole cells or from nuclear or cytoplasmic fractions was extracted using TRIzol reagent (Invitrogen; Thermo Fisher Scientific, Inc.) and treated with DNase I (Promega, Madison, WI, USA). cDNA was reverse transcribed from 500 ng of total RNA with hexamers using a PrimeScript RT reagent kit (Takara Bio, Inc., Kusatsu, Japan). qPCR was then performed in a 25 μl reaction using TB Green Premix Ex Taq II reagent (Tli RNaseH Plus; Takara Bio, Inc.) on an Applied Biosystems 7500 Real-Time PCR System. The thermocycler protocol was as follows: 95°C for 30 s followed by 40 cycles of 95°C for 10 s and 60°C for 30 s. The primer sequences are listed in (Supplementary information, Table S12). The Cyto/Nuc ratio was calculated using ΔCq values. All experiments were performed in triplicate.

### CRISPR/CAS9 correction

Biallelic mutations in MAD-iPSCs (LMNA^R527C/R527C^) were corrected via previously described methods^63, 64^. Briefly, we utilized the electroporation of CAS9 protein and gRNA-tracrRNA, along with a single stranded oligonucleotide (ssODNA) donor template. The gRNA and donor template sequences are listed in (Supplementary information, Table S13). Briefly, the gRNA complex was first prepared by incubating equal volumes of 100 µM of crRNA and TracrRNA-ATTO-550 (ALTR-HDR, IDTA) at 95°C for 5 min, followed by incubation at room temperature for 20 min. The RNP complex was made by adding 2 µl sp.Cas9 Nuclease V3 protein (61 µM or 10 µg/µl; IDTA) with 2.5 µl gRNA (tracrRNA+crRNA) and incubating at RT for 20 min. Simultaneously, IPSCs were incubated with 5 µm of RockIn (STEMCELL Technologies, Inc.) for 2-3 h before dissociating with Accutase. Cells were washed with PBS, and 1×10^6^ cells were centrifuged at 300 x *g* for 5 min. Cells were mixed with complete p3 electroporation solution (from a Lonza 4D nucleofection kit; Lonza Group AG, Basel, Switzerland) followed by addition of the RNP complex and 500 pmoles of donor ssODNA (Alt-R HDR; IDTA). Electroporation with the CA-137 program was performed 3-5 min after this addition (Lonza Group AG). Following this, cells were quickly but gently resuspended in six-well coated plates with Synthemax (Sigma-Aldrich; Merck KGaA) in MtesR-plus medium supplemented with CloneR (STEMCELL Technologies, Inc.), and plates were incubated for 48 h at 32°C with 5% CO2. After 48 h, cells were maintained at 37°C. IPSCs were dissociated with Accutase once again, and passed through a single cell filter before being seeded at a density of 500 cells in a 10 cm dish. When single IPSC colonies began to appear after ∼10-14 days, these were manually selected and seeded in 96 well plates, in replicate.

### Mitochondrial and cytoplasmic calcium quantification

Mitochondrial Ca^+2^ and cytoplasmic Ca^+2^ were measured using two different dyes: Rhod-2,AM (Yeasen Biotechnology Co., Ltd., Shanghai, China; cat. no. 40776ES50) and Fluo-4,AM(Yeasen Biotechnology Co., Ltd.; cat. no. 40704ES50).

Cells were incubated with 0.2 µM Rhod-2,AM (37°C, 20min) to measure mitochondrial Ca^+2^ levels and with 2 µM Fluo-4,AM (37°C, 20 min) to measure intracellular calcium levels. This was followed by washing with PBS and incubation for 20 min in PBS, and live imaging with Ex/Em of 488/525 and 550/585 for Fluo-4 and Rhod-2,AM-stained cells, respectively. Mitochondrial Ca^2+^ was blocked with CGP37157 at 30 µM (37°C, 20 min). Quantitative analysis was performed using Image J software (ImageJ 1.8.0;National Institutes of Health, Bethesda, MD, USA).

### Chromatin extraction

iMSCs were lysed in hypotonic lysis buffer (10 mM Tris-HCl pH 8.0, 1 mM KCl, 1.5 mM MgCl2 and 1 mM DTT) containing protease inhibitors, and the intact nuclei were pelleted by centrifugation at 14000 x g at 4°C for 10 min. Cell supernatant was discarded, and the nuclei were re-suspended in 400 µl 0.2 M sulfuric acid and incubated at 4°C for 30 min. The samples were again collected by centrifugation at 14000 x g at 4 °C for 10 min, and the supernatant containing the histones was collected. Trichloroacetic acid was added to the histones to a final concentration of 33%, and the samples were incubated on ice for 30 min. Following this, the samples were centrifuged at 14000 x g at 4°C for 10 min, and the histone pellets were collected, washed with acetone, and dissolved in ddH2O.

### Treatment with IL-6 and/or IL-6 inhibitors

Stat3 was activated in healthy iMSCs by adding the cytokine IL-6 (PeproTech, Rocky Hill, NJ, US; cat. no. 200-06-5UG) into the culture medium at the 20 ng/ml concentration for (24 hours in 37 °C). Cells were then treated with 10 µg/ml tocilizumab (MedChemExpress LLC; cat. no. HY-P9917) or 0.25 µm tofacitinib (MedChemExpress LLC, cat. no.HY-40354) for 24 hours 37°C. To confirm the effects of LMT-28, a synthetic IL-6 inhibitor, cells were initially starved for 24 h (only MSC basal medium, without supplementation) and then treated were with 30 µm LMT-28 (Medchem Express LLC; cat. no. HY-102084) for 1 h,37 °C. Mitochondrial membrane potential was also rescued by treating cells with the mTOR inhibitor AZD-8055 (MedChem Express LLC; cat. no. HY-10422) at concentration of 500 nM for 24 h, 37°C.

### Antibodies

Details of antibodies used for iMSC surface antigen identification are presented in Table 1. For western blot analysis, details of primary antibodies used are presented in Table 2. The following secondary antibodies used in western blot analysis were purchased from Cell Signaling Technology (Danvers, MA, USA) and diluted at 1:5000: HRP-linked rabbit IgG antibodies (cat. No. 7074S) and HRP-linked rabbit IgG antibodies (cat.no. 7076S).

Details of antibodies used for immunofluorescence are presented in Table 3. The following fluorescence-conjugated secondary antibodies were purchased from Cell Signaling Technology and diluted at 1:800: rabbit Alexa Fluor-488 antibodies (cat. no. 2975S), rabbit Alexa Fluor-594 antibodies (cat. no. 8760S), mouse Alexa Fluor-594 antibodies (cat. no. 8527S) and mouse Alexa Fluor-488 antibodies (cat. no. 4408S). In addition, DAPI (1:5000; cat. No. D9542; Sigma-Aldrich; Merck KGaA, Darmstadt, Germany) was used for nucleic acid staining, and phalloidin-iFluor 488 (1:1000; cat. no. ab176753; Abcam, Cambridge, UK) and phalloidin-iFluor 594(1:1000; cat. no. ab176757; Abcam) were used for binding to actin filaments.

### Statistical analysis

Statistical analyses were performed using GraphPad Prism Software Version 9.4 (GraphPad Software, Inc., La Jolla, CA, USA). Comparisons were made using two-tailed unpaired Student’s t-tests and one-way ANOVA followed by Student-Newman Keuls post hoc tests. In-case of multiple comparisons, Two-way analysis of variance with Turkey’s multiple comparison was utilized. P<0.05 was considered to indicate a statistically significant difference.

## Acknowledgement

This work is supported, in part, by National Natural Science Foundation of China (2072480、32060157 and 32100603), and Shenzhen Commission of Development Reform (Funding for Shenzhen Engineering Laboratory for Orthopedic Diseases and Regenerative Technologies). The authors would like to thank Dr. Jessica Tamanini (Shenzhen University and ETediting).

## Authors Contribution

Conceptualization: AAP, GZ; Funding acquisition: GZ, WS and YZ; Writing original Draft: AAP and XY; Review and Editing: GZ, SZ, TW, GW, XZ, MZ; Mouse model: GA, AF, AAP and LH; Exosomes purification: AAP, AF, XY and LH. MSCs differentiation: JL, ZL and XY. IPSCs generation and characterization: TW, JL, AAP, ZL. Assessment of patients PBMCs markers clinical data: WS, LZ, MZ, TL, XZ, GW and GZ. Methylation, acetylation status was evaluated by CAA, XY and AAP. CRISPR/CAS9 correction: AAP, JL and XY. Unmentioned contribution to the manuscript was carried by AAP and/or XY. All authors have approved the submitted version of the manuscript.

