## supplementary information for "MAM-STAT3-induced upregulation of mitochondrial Ca^+2^ causes immunosenescence in patients with type A mandibuloacral dysplasia"

Running Title: Cell senescence and inflammaging in progeroid syndrome.

**SUPPLEMENTARY NOTE 1: Patients’ clinical features**

4 patients from three different families were brought to the first affiliated hospital of Guangxi medical university from January 2017 to December 2018. All of them showed hallmarks of severe atypical progeroid symptoms and scleroderma-like skin changes. Three patients of two family members had homozygous LMNA p.R527C mutation and denoted as MAD1, MAD2 and MAD3. Whereas one patient has heterozygous LMNA G608G mutation and denoted here as HGPS1. Except for the patients, other members of the family have no obvious abnormal symptoms. No obvious genetic history was present in the family, and parents had denied the consanguineous marriage. The clinical characterization and radiological observations are described in Fig.1, Fig.S1 and Fig.S2. All of them suffered from growth retardation,	“bird like” facial characterization and horse-riding stance. Large head, sparse hairs, beak-shaped nose, protruding eyes and severe mandibular and tooth hypoplasia was also in common. Along with skeletal hypoplasia micrognathia, bilateral clavicle hypoplasia, short rod-shaped finger/toe end, severe deformity of the proximal knuckle joints of both hands, and a bilateral distal clavicle hypoplasia. These patients also showed less subcutaneous fat on the extremities of the trunk but more subcutaneous fat on the cheeks and neck. Patient skin showed scleroses-like changes (hyperplasia of dermal collagen, fibrosis, and inflammatory cell infiltration). Patients with p.R527C mutation also showed elevated platelets and decreased serum creatinine. Creatine kinase isoenzyme (CK-MB) and lactate dehydrogenase (LDH) were slightly elevated in MAD patients and only LDH in HGPS patient. Notably, Liver and kidney function was normal, and all patients presented with normal hearing and cognitive function.

MAD1:

The MAD1 patient (female, age:3) belongs to the family 1, which came from Liuzhou city, Miao Nationality, Guangxi Zhuang autonomous region, China. She was admitted to hospital for suffering skin abnormality and was diagnosed with a moderate scleroderma when she was 3 years old. At the age of 3 y, she had a height of 82cm (<P1), weight of 8 Kg (<P1), head circumference of 44.5 cm (<P1), she was the second child in family that delivered vaginally. The first abnormal manifestation was found when she was 6 months old. The skin biopsy features were suggestive of proliferated dermal collagen fibers and fibrosis. Embryonic-like adipocytes were also infiltrated in the subcutaneous fat. She showed classical features of acroosteolysis of hands, feet with knuckles were absorbed, shortened or partially absent. Mandibular recession and distal clavicular hypoplasia along with subcutaneous lipoatrophy can be observed, whereas increased subcutaneous fat is observed in localized region, especially in the cheeks.

MAD2:

The MAD2 patient (male, age: 5) belongs to family 2 which came from Hechi city of Gangxi Zhuang autonomous region of Zhuang nationality. His abnormal symptoms started to manifest by the age of 1 year, when he began to show progressive loss of hair and eyebrow, itchy skin, along with the increased in skin pigmentation. At the age of 5 years his height was 86 cm (<P1), weight 8 kg (<P1). He was the second child in the family that born with cesarean section. Radiographs in the Fig. S1 showed the acroosteolysis with brachydactyly, which is more pronounced distally. Additionally, clavicular hypoplasia, loss of partial bone, calvaria-mandibular disproportion and pear-shaped breast is also noted. H&E staining from the left forearm skin biopsy showed that the dermal collagen fibers were proliferated and dense. Infiltration of lymphocytes and plasma cells, collagen fibers are thick with the degeneration of hyaline Fig S2

MAD3:

The MAD3 patient (male, ag**e: 7)** also belongs to family 1 and the elder brother of MAD1 patient. At the age of 7 years his height was 90 cm (<P1), weight 9.5 kg (<P1) and head circumference 46.5 cm (<P1), he was the first child of his parents that had delivered vaginally. His clinical manifestations are basically the same, but more severe than MAD1. Skin sclerosis and subcutaneous fat atrophy of the limbs are more obvious, and the lesions in both fingers are more serious (all knuckles are absorbed, shortened, and partially absent), with severe scoliosis (Fig. 1C). The patient suffered from skin itching at the age of 1 yr, accompanied by developing hair loss and darkening of the skin on the back of both hands. Around 2 years old, he presented with swelling of fingers, skin sclerosis, decreased joint motion and progressive flexion deformity of the proximal knuckle joints of both hands was observed, which was diagnosed as severe scleroderma. The skin biopsy features were suggestive of dermal collagen fibers and reduction of hair follicles. Infiltration of embryo-like cells was also noted with higher frequency (Fig. S2B1-2).

HGPS1:

The HGPS1 patient (male, 1 y) belong to family 3 that came from Beihai City, Guangxi Province. Patient presented at the age of 9 month to first affiliated hospital of Guangxi medical university due sparse hair and swelling of the lower limbs. The swelling of the limb was started at the age of 3 month, without any other discomforts. Child was born through vaginal delivery and parents denied for consensual marriage. The first pregnancy was aborted for which the details were not disclosed. Growth retardation, large head, prominent forehead, sunken eye sockets, pointed nose, obvious exposure of scalp and trunk veins, hardening of the skin on the abdomen and lower limbs, with no special changes in the upper arms. HGPS patients showed less subcutaneous fat throughout the body (generalized lipodystrophy), compared to MAD patients, though the HGPS patient's skeletal dysplasia is less severe. The result of skin biopsies was presented in Supplementary Fig. S2 C1-C2. At the age of 1 years his height was 68 cm (<P1), weight 7kg (<P1) and head circumference 45.5 cm, he was the second child of first birth, vaginal delivery.

**Supplementary Fig S1: Radiographical features of MAD2 patients.**

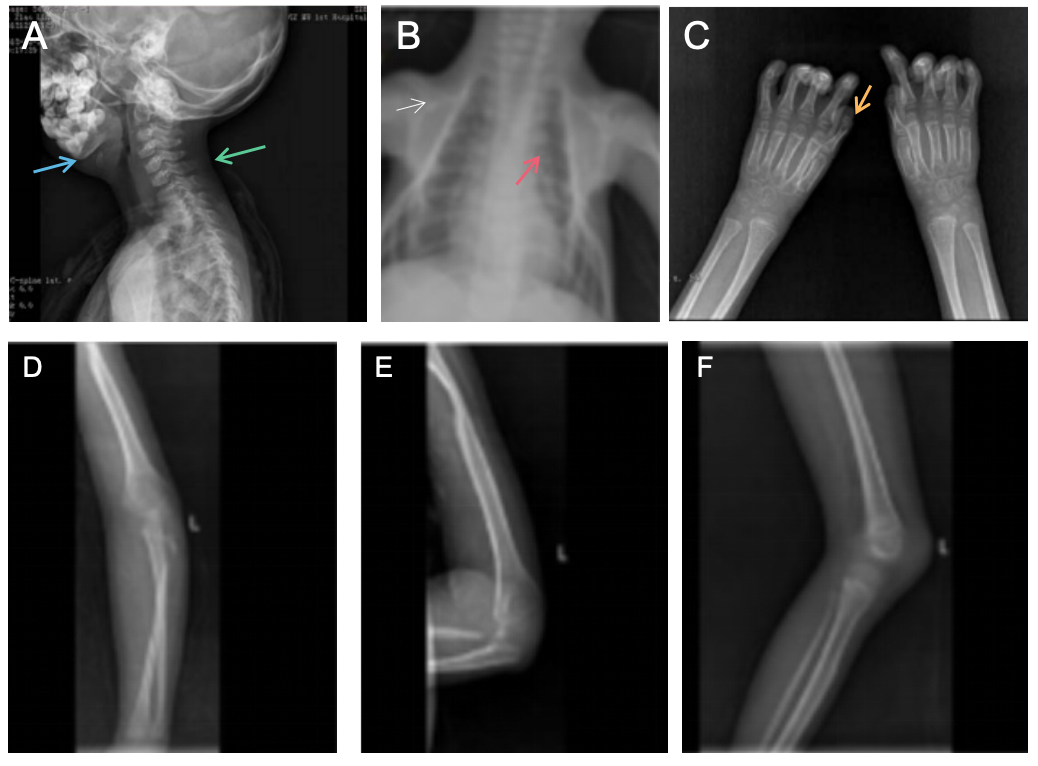

A. Lateral neck radiograph showing micromandible (blue arrow), calvaria-mandibular disproportion, and right sternocleidomastoid dysplasia resulting in torticollis (green arrow).

B. X-ray showing narrow and pear-shaped chest (red arrow) and severe bilateral or dissolved clavicular hypoplasia (white arrow).

C. Hand radiographs of patient showing short, bone defect and club-shaped distal phalanges of all digits (yellow arrow).

D-F. Four extremities radiographs of patient showing decreased bone density in the extremities and loss of partial bone.

**Supplementary Fig S2. Haematoxylin and eosin staining of skin biopsies**

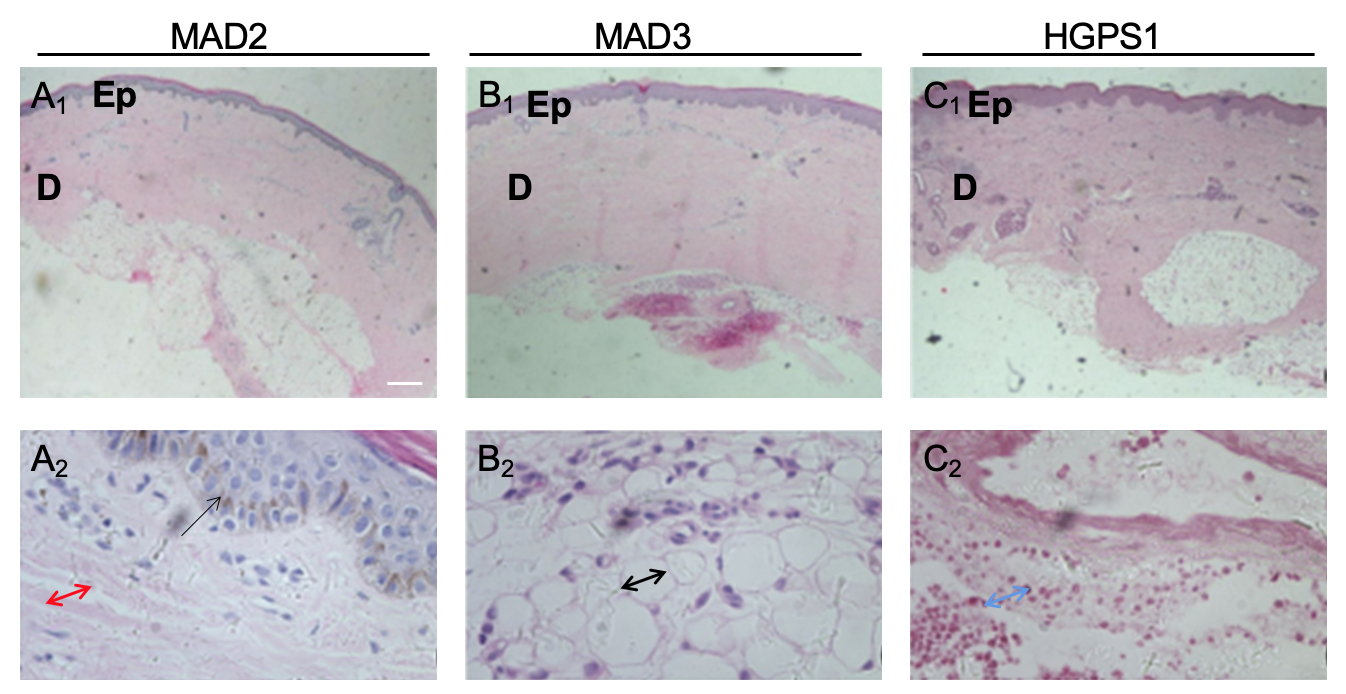

A_1_. HE-staining of the left forearm showed that the dermal collagen fibers were proliferated and dense.

A_2_. Infiltration of lymphocytes and plasma cells (black arrow)，collagen fibers are thick and hyaline degeneration (double headed red arrow).

B_1_. In abdominal skin, we observed hyperplasia of dermal collagen fibers and reduced hair follicles.

B_2_. Black double headed arrow represents embryo-like cells.

C_1_. A normal epidermis, hyperplasia of dermal collagen, fibrosis and hyalinization.

C_2_. Fibrotic degeneration of patient’s right calf intradermal blood vessel wall. The vessel wall and surrounding areas also showed more eosinophils (blue double headed arrow). Scale bar: upper panel = 1000 μm, lower panel = 100 μm

**Supplementary Fig S3. Identification of pluripotency markers in iPSCs.**

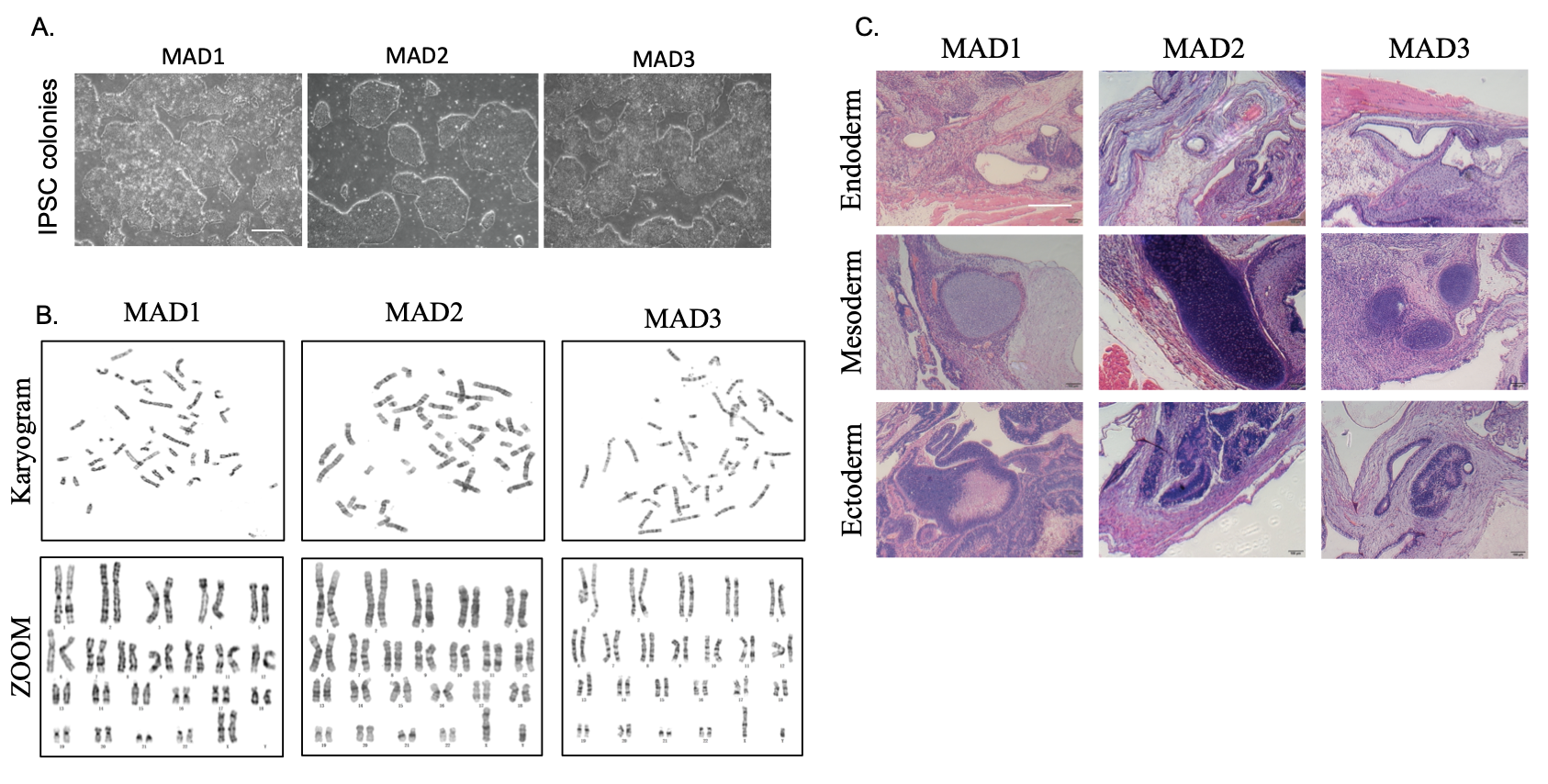

A) Representative images of PBMC-derived induced pluripotent stem cell colonies, scale bar = 200μm. B) Karyotype analysis revealed the normal karyotypes of 46 chromosomes of MAD1, MAD2 or MAD3 patient PBMC derived iPSCs. C) A representative microscopic image of histological section is stained with H&E of teratoma derived from MAD-iPSCs. Arrows point to areas of interest: Ectoderm, neural tissue; Mesoderm: cartilage tissue; Endoderm, intestinal epithelial tissue. Scale bar = 100μm.

**Supplementary Fig S4. Confirmation of pluripotency by IPSCs markers.**

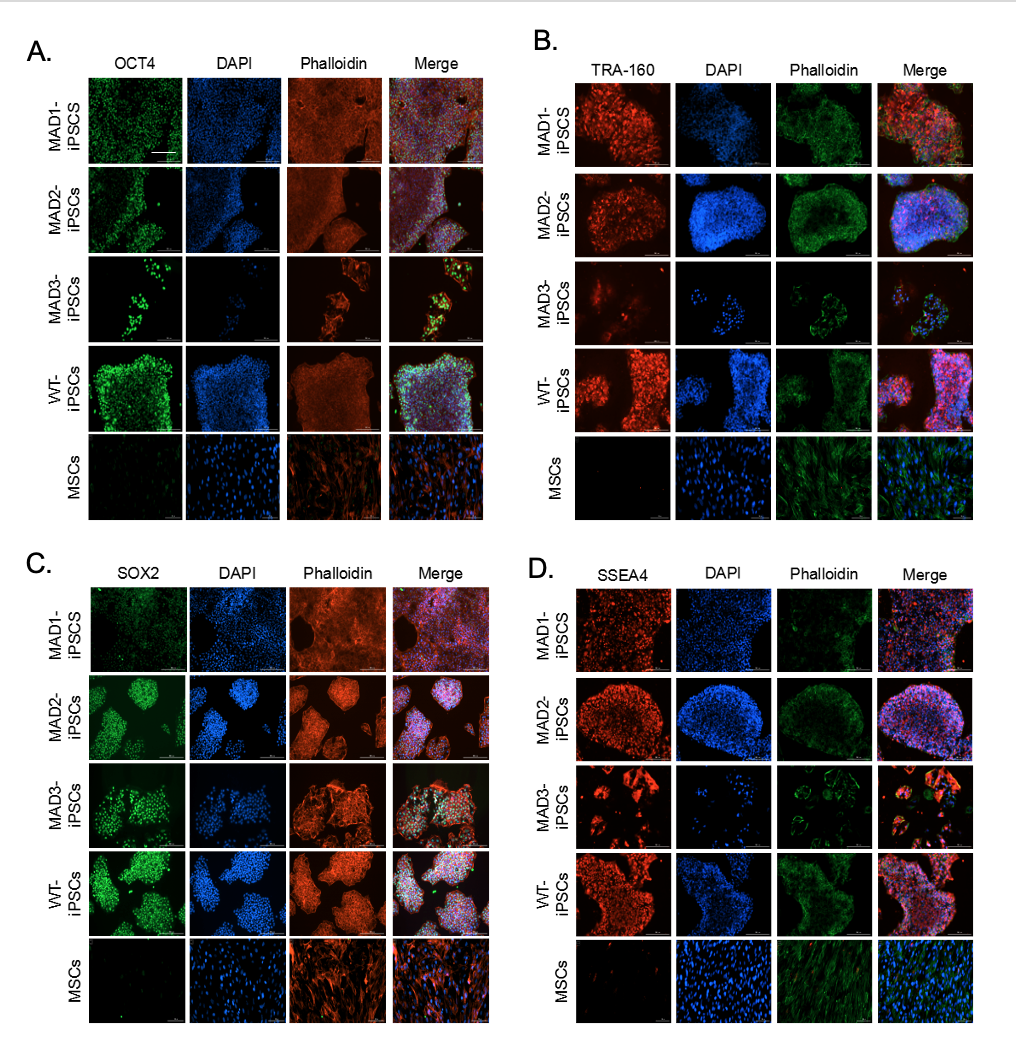

A-D) Immunofluorescence staining OCT4 (green), TRA-160 (red), sox2 (green), SSEA4 (red); DAPI (Blue); Phalloidin dye is used here for counter staining. Mesenchymal stem cells (MSCs) were taken as negative control. Scale bar = 200 µm.

**Supplementary Fig S5. iPSCs marker confirmation after 40 passages**

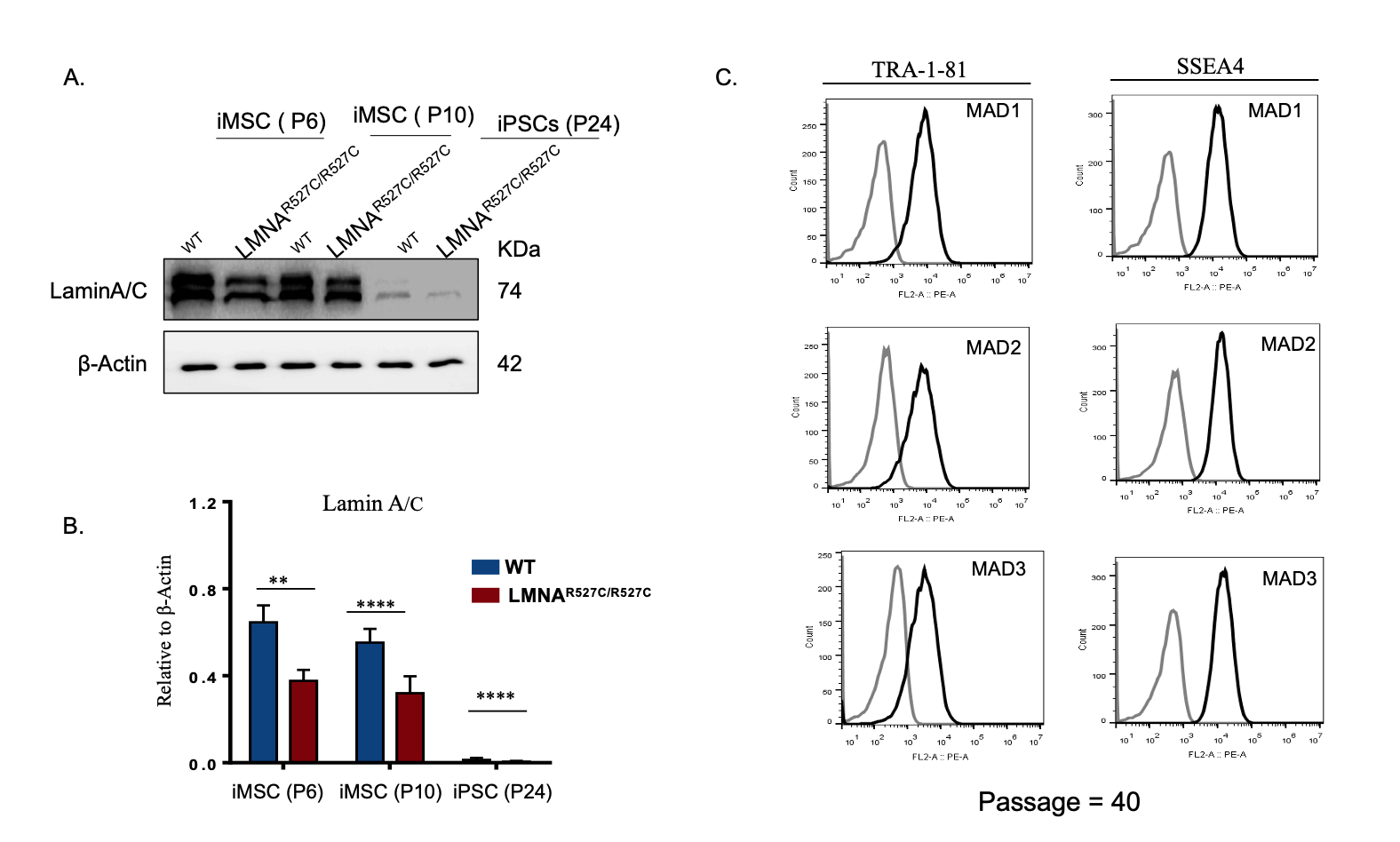

A, B) Western blot of LMN A/C on two different passages wild type (WT) and MAD iMSCs along with the cell lysate WT and MAD iPSCs. At any given passage WT cells expressed more LMN A/C compared to MAD, though LMN A/C expression increased with the increase in passage number. Here, iPSCs also showed hairline band suggesting the presence of LMN A/C in our iPSCs. No progerin or pre-laminA expression was detected on MAD derived iMSCs. C) Flowcytometry of MAD patient derived iPSCs at passage 40, showed presence of pluripotent marker TRA-1-81 and SSEA4.

**Supplementary Fig S6. Characterization of iPSCs derived iMSCs.**

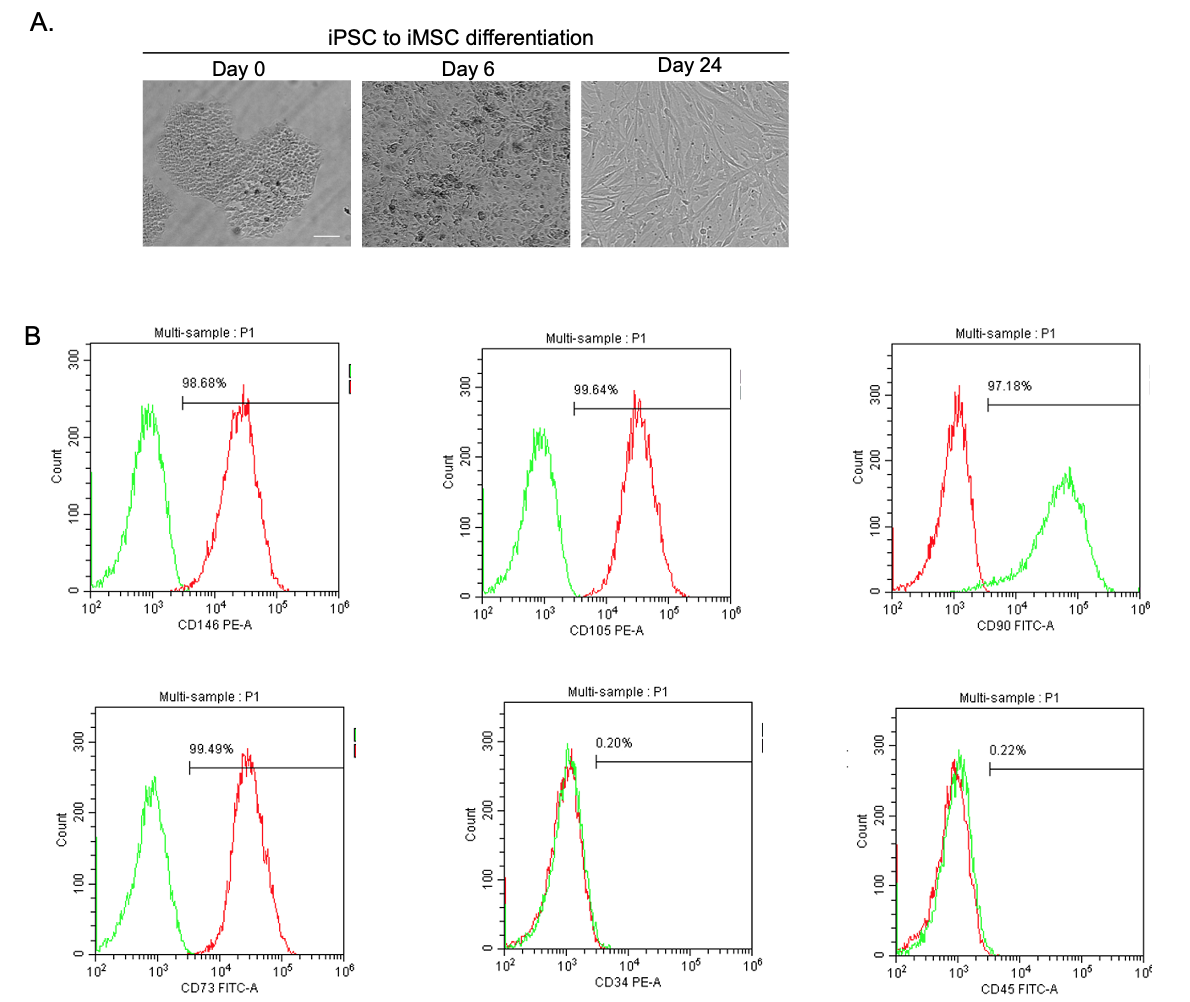

A. Image of long-term cultured iPSC clone before differentiation (i) intermediate phase of induced iPSCs (ii), and iMSCs (iii), scale bar = 200 μm.

B. Flow cytometry analysis shows the positive to iMSCs markers: CD146 (98.68%), CD105 (99.64%), CD90 (97.18%), CD73 (99.49%); and negative to markers CD34 or CD45. n=3.

**Supplementary Fig S7. CRISPR-Cas9-Mediated LMNAp.R527C Correction**

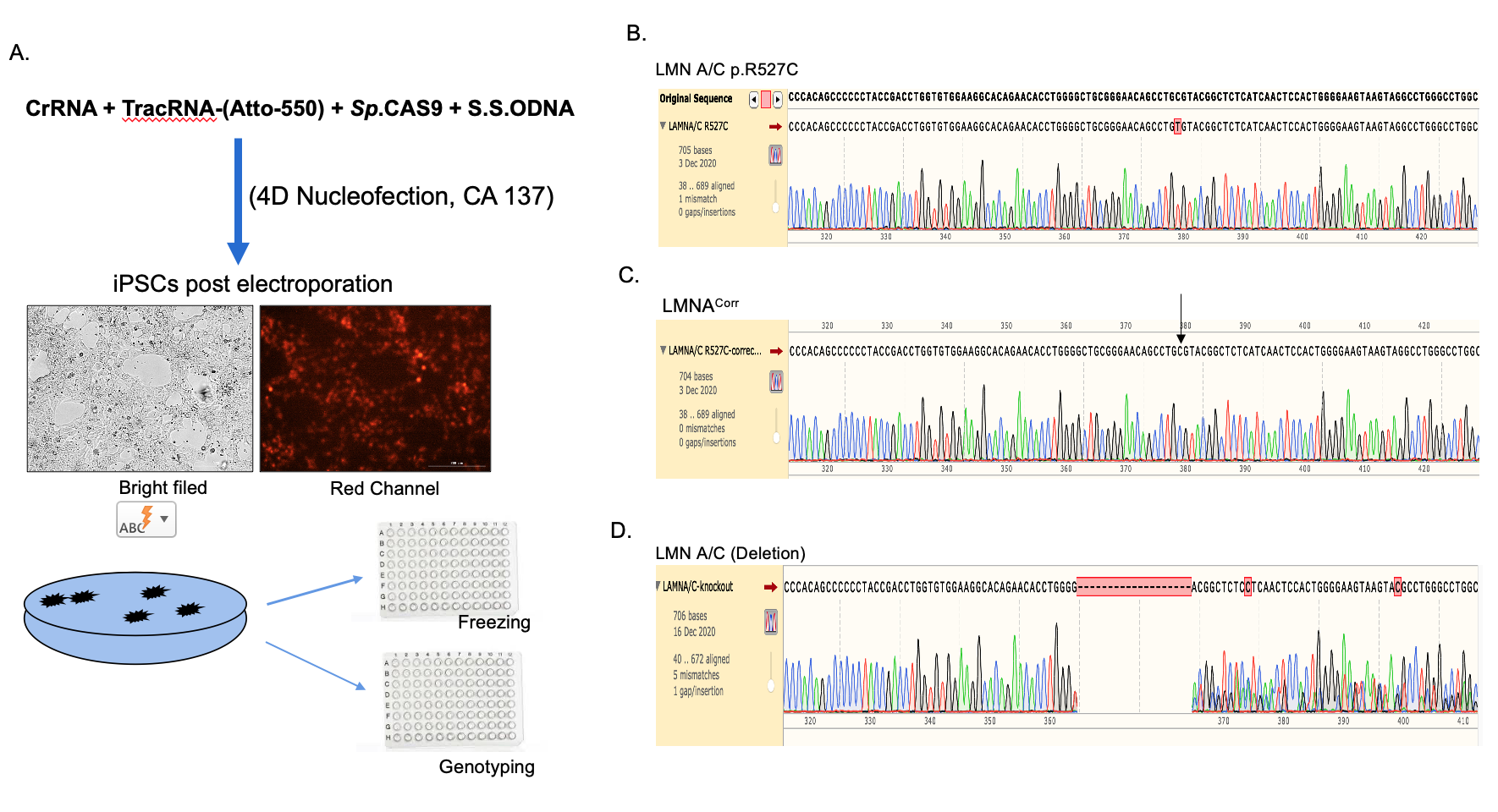

A) RNP complex was electroporated using 4D nucleofector followed by seeding the cells. Nucleofection efficiency was ~99%, as shown here on red channel. After 4 days for electroporation, few hundred (300 to 500) cells were seeded on 10 cm^2^ petri dish until the individual iPSC colonies were formed. Colonies were manually picked and divide them in two 96 well plate. Scale bar = 200 µm. B) un-edited MAD-iMSCs cells after electroporation C) homozygous corrected cells. In our setup we able achieve 23% of total homozygous correction. D) Around 50% of the iPSC clones had the deletion.

**Supplementary Fig S8: Rectification of Nuclear abnormality and senescence marker in MAD cells after correction of mutation with CRISPR/CAS9**

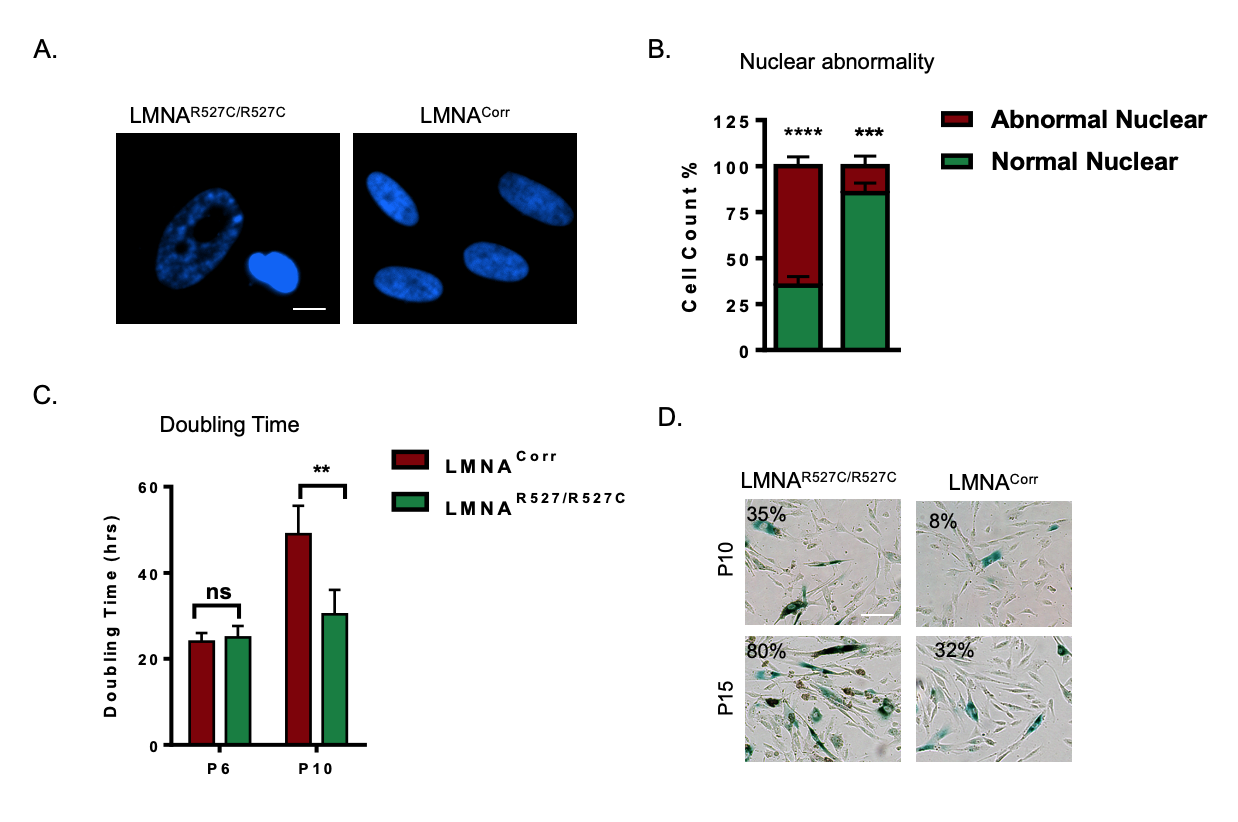

A-B) Fluorescence images of nuclei stained with DAPI (blue) showed the rectification nuclear blebbing and honeycomb nuclei in the LMNA^corr^ iMSCs. The scale bar represents 20 µm (n=8). (C) Cell proliferation plotted against cell doubling time. D) Cell senescence was quantified at the indicated passage in LMNA^R527C/R527C^ and LMNA^corr^ iMSCs with beta-galactosidase staining**.** *p<0.05, **p<0.01, and ***p<0.001, with comparisons indicated by lines. Scale bar = 100 µm.

**Supplementary Fig S9. Screening the retention of mitochondrial membrane transcripts in the nucleus of LMNA p.R527C cells.**

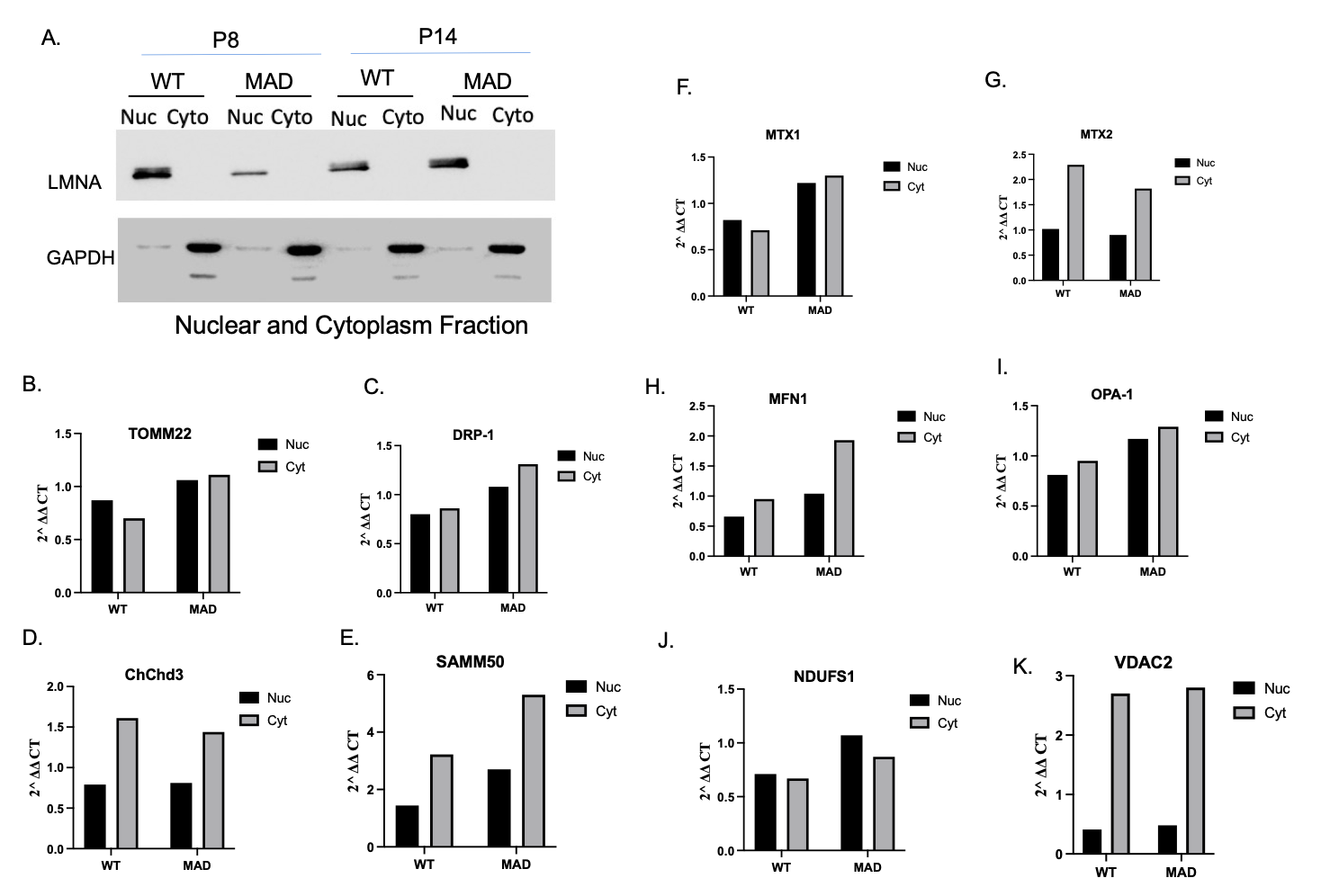

A) Western blot of nuclear and cytoplasmic fraction of WT and MAD iMSCs of two different passages. B-K) Quantitative real time PCR of different outer and inner mitochondrial transcripts expressed as 2^–∆∆Ct^ values. Though some of transcripts for instance, ChChd3 expression is higher in WT cytoplasm or SAMM50 has increased expression in MAD patient, but the there was no significant difference in the association of nuclear retention of integral mitochondrial membrane transcripts with the p.R527C LMNA mutation.

**Supplementary Fig S10:**

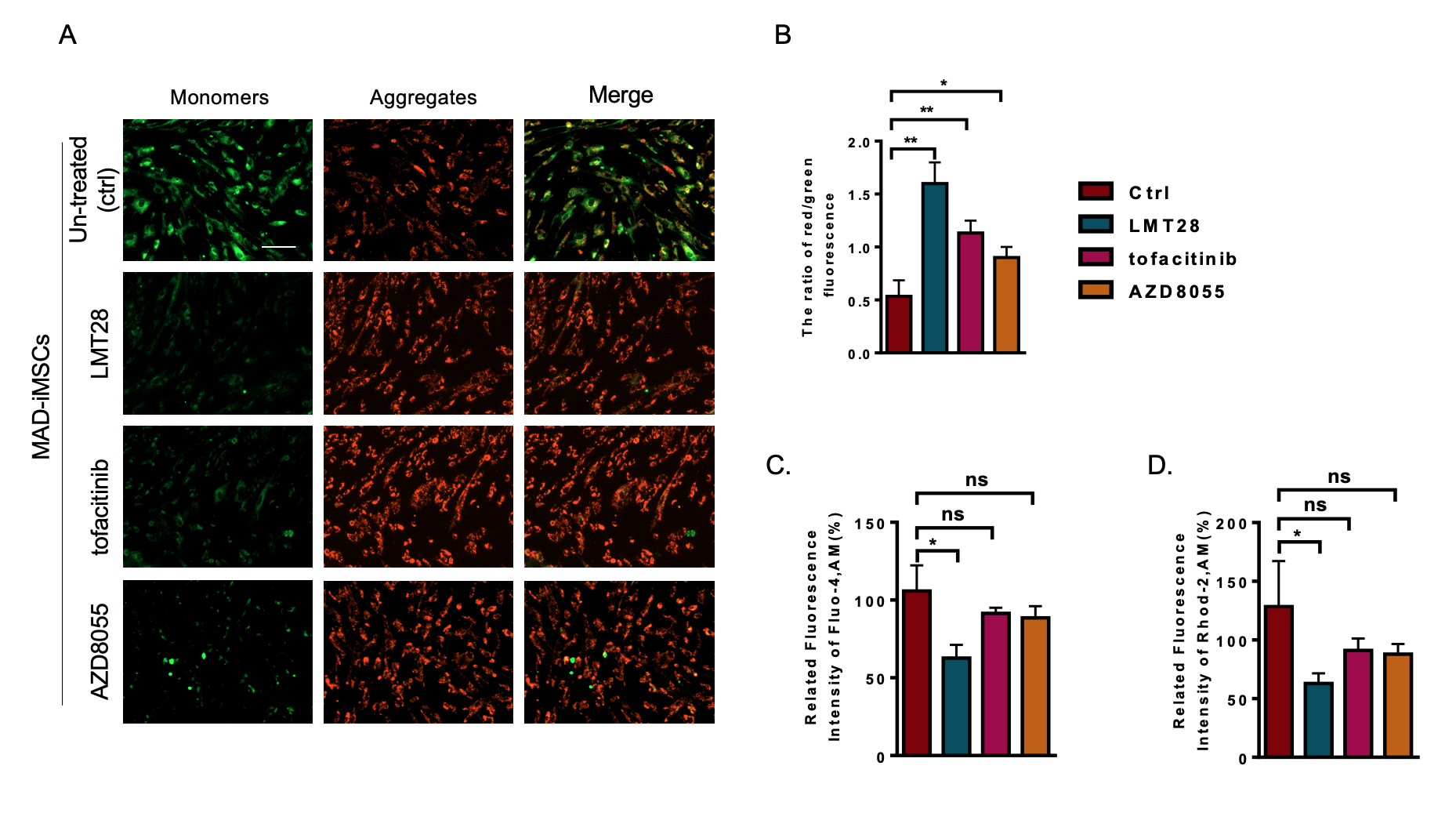

**A and B)** MAD-iMSC cells at passage 10 were treated with 30 uM of LMT-28, 0.25 uM of Tofacitinib and 500 nM AZD8055 for the indicated duration in method section. JC-1 staining revealed that cells treated with direct and indirect IL-6 inhibitors regained their lost mitochondrial membrane potential. Scale bar = 200 µm. **(C and D)** Graphs showed the quantification of Mitochondrial and cytoplasmic Ca^+2^ levels that was observed with Rhod-2,AM and Fluo-4,AM dyes at Ex/Em 4550/585 and 488/425 nm, respectively.

**Supplementary Fig S11**

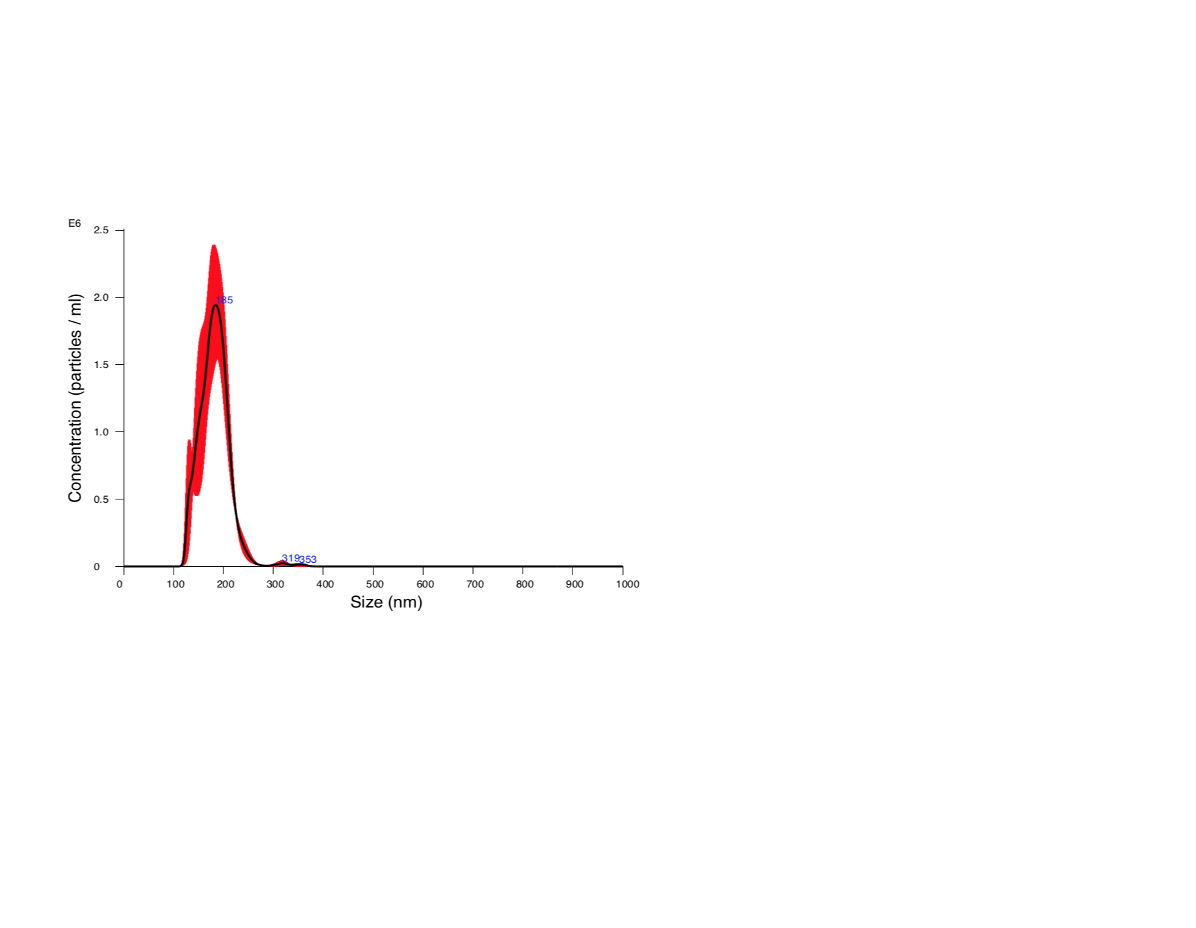

Nanoparticle tracking analysis showed that mostly MAD-iMSc-EVs were 180nm in diameter.

**Supplementary Fig. S12**

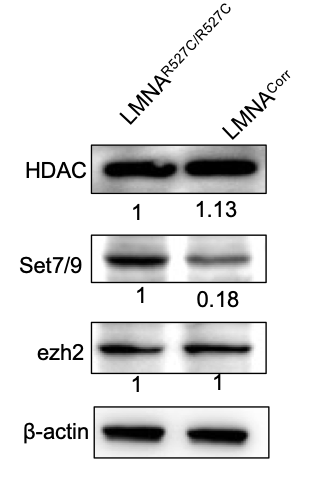

Western blot analysis of MAD-iMSC showed the various expression of enzyme that controls the heterochromatin modification. Set7/9 expression was found to be higher in LMNA p.R527C mutated cells compared to corrected or wild-type control

**Supplementary Tables**

**Table S1: Screening for Autoimmune antibodies in MAD patients**

| Antibody | MAD1 | MAD2 | MAD3 | C1 | C2 | C3 | Normal Range |
| --- | --- | --- | --- | --- | --- | --- | --- |
| Anti-Sm | <3.7 | <3.7 | <3.7 | <3.7 | <3.7 | <3.7 | 0-25 AU/ml |
| Anti-nuclear | Neg | Neg | Neg | Neg | Neg | Neg | Neg (1:80) |

Antinuclear antibodies were quantified using indirect immunofluorescence from plasma samples. Ratio of 1:80 was the cut-off negative titre for ANA.
Patients: MAD1 (Female, 3 y); MAD2 (Male, 5 y), MAD3 (Male, 7 y)
Healthy Controls: C1 (Female, 3 y); C2 (Male, 5 y); C3 (Male, 7 y)

**Table S2: Quantification of cytokines from serum samples**

| **pg/mL** | **C1** | **MAD1** | **C2** | **MAD2** | **C3** | **MAD3** | **C4** | **HGPS** | **C5** | **C6** | **C7** |
| --- | --- | --- | --- | --- | --- | --- | --- | --- | --- | --- | --- |
| **IL-5** | 4.36 | 1.74 | 3.40 | 3.64 | 1.92 | 3.17 | 8.29 | 3.90 | 5.23 | 1.94 | 1.73 |
| **IL-13** | 2.30 | 2.12 | 3.69 | 6.02 | 1.90 | 4.77 | 6.12 | 8.01 | 4.50 | 2.38 | 2.04 |
| **IL-2** | 2.04 | 2.57 | 2.16 | 4.15 | 2.38 | 4.15 | 4.74 | 4.66 | 7.95 | 1.73 | 1.57 |
| **IL-6** | 7.16 | 12.92 | 3.30 | 28.27 | 5.00 | 16.79 | 20.55 | 15.89 | 49.85 | 7.79 | 3.74 |
| **IL-9** | 5.66 | 24.34 | 1.70 | 52.38 | 2.71 | 24.86 | 15.75 | 13.74 | 13.03 | 7.41 | 2.31 |
| **IL-10** | 2.60 | 6.53 | 1.37 | 11.30 | 1.61 | 7.86 | 11.58 | 3.60 | 5.13 | 2.69 | 1.35 |
| **IFN-γ** | 14.66 | 5.67 | 34.28 | 16.98 | 25.04 | 20.95 | 87.33 | 230.60 | 39.37 | 20.01 | 4.37 |
| **TNF-ɑ** | 4.75 | 2.45 | 6.46 | 3.27 | 5.48 | 3.49 | 7.58 | 15.64 | 4.60 | 4.96 | 2.37 |
| **IL-17A** | <1.09 | <1.09 | <1.09 | <1.09 | <1.09 | <1.09 | <0.47 | 1.34 | <1.09 | <1.09 | <1.09 |
| **IL-17F** | 1.54 | 0.85 | 2.35 | 1.72 | 0.96 | 1.42 | 1.62 | 7.48 | 0.91 | 1.07 | 1.13 |
| **IL-4** | 1.67 | 1.66 | 2.34 | 5.16 | 1.88 | 3.93 | 5.05 | 12.35 | 1.84 | 1.84 | 1.43 |
| **IL-21** | 1.41 | 1.30 | 2.29 | 2.24 | 2.47 | 2.15 | 9.64 | 7.34 | 2.62 | 2.47 | 1.72 |
| **IL-22** | 1.64 | 1.23 | 1.75 | 2.46 | 1.72 | 4.70 | 11.60 | 4.03 | 1.19 | 1.27 | 1.39 |

Patients: MAD1 (Female, 3 y); MAD2 (Male, 5 y), MAD3 (Male, 7 y)
Controls: C1 (Female, 3 y); C2 (Male, 5 y); C3 (Male, 7 y); C4 (Male, 1 y), C5 (Female, 65 y); C6 (Male, 65 y); C7 (Male, 65 y)

**Table S3: Sub-set of T-lymphocyte markers**

| **Cell-Type** | **Marker** | **C1** | **MAD1** | **C2** | **MAD2** | **C3** | **MAD3** | **C4** | **HGPS** | **C5** | **C6** | **C7** |
| --- | --- | --- | --- | --- | --- | --- | --- | --- | --- | --- | --- | --- |
| **Total T lymphocytes** | CD3^+^ | 61.8 | 67.70 | 65.2 | 51.11 | 59.6 | 61.05 | 54.74 | 65.4 | 47.77 | 42.17 | 51.94 |
| **Helper T cells** | CD3^+^ CD4^+^ | 58.6 | 50.63 | 43.5 | 46.59 | 58.4 | 44.15 | 48.83 | 71.6 | 74.45 | 46.79 | 47.41 |
| **Killer T cells** | CD3+CD8+ | 35.7 | 38.03 | 29.4 | 39.56 | 31.1 | 42.37 | 44.32 | 23.8 | 24.55 | 50.86 | 49.20 |
| **Th/killer T cells ratio** | | 1.64 | 1.33 | 1.48 | 1.18 | 1.87 | 1.04 | 3.03 | 1.10 | 0.92 | 3.03 | 0.92 |
| **Double +ve T-cells** | CD3^+^(CD4^+^CD8^+^) | 0.38 | 0.18 | 0.22 | 0.64 | 0.24 | 0.38 | 0.53 | 0.28 | 1.01 | 1.15 | 1.36 |
| **Activated-T cells** | CD3^+^CD25 | 7.41 | 6.51 | 6.29 | 16.44 | 7.57 | 8.08 | 5.94 | 9.29 |  | 13.31 | 10.71 |
| **Activated Helper T cells** | (CD3^+^CD4^+^) CD25^+^ | 12.4 | 12.44 | 15.3 | 36.20 | 12.4 | 14.97 | 12.01 | 12.9 |  | 26.94 | 21.92 |
| **Activated-Killer T cells** | (CD3^+^CD8^+^) CD25^+^ | 0.16 | 0.14 | 0.38 | 0.63 | 0.29 | 0.28 | 1.11 | 0.54 |  | 1.13 | 0.43 |
| **γ δ T cells** | CD3+TCRγ δ + | 4.83 | 11.31 | 31.8 | 17.81 | 11.8 | 12.74 | 6.66 | 4.38 | 2.00 | 4.51 | 7.49 |
| **Activated γ δ T cells** | CD3+(γ δ T+CD25+) | 0 | 0.03 | 0.80 | 0.60 | 0.37 | 1.13 | 0.15 | 0.15 | 0.04 | 0.18 | 0.28 |
| **Senescent T cells** | CD3+CD57+ | 3.56 | 9.82 | 4.96 | 7.64 | 6.85 | 16.27 | 13.23 | 0.29 | 13.05 | 27.09 | 36.05 |

Peripheral blood (subset of T cells): Total lymphocyte count or the subset of T cells of MAD patients were not varied with their respected controls. Only the percentage of senescent cell population have higher number

Patients: MAD1 (Female, 3 y); MAD2 (Male, 5 y), MAD3 (Male, 7 y)
Controls: C1 (Female, 3 y); C2 (Male, 5 y); C3 (Male, 7 y); C4 (Male, 1 y), C5 (Female, 65 y); C6 (Male, 65 y); C7 (Male, 65 y)

**Table S4: CD3^+^ T cells senescent marker population.**

| **Phenotype (%)** | **C1** | **MAD1** | **C2** | **MAD2** | **C3** | **MAD3** | **HGPS** | **C5** | **C6** | **C7** |
| --- | --- | --- | --- | --- | --- | --- | --- | --- | --- | --- |
| **CD3+CD57^+^** | 3.88 | 14.69 | 4.31 | 11.14 | 6.51 | 23.75 | 2.01 | 14.92 | 31.31 | 45.01 |
| **CD3+KLRG -1^+^** | 6.13 | 25.60 | 30.54 | 63.18 | 15.73 | 40.67 | 7.07 | 19.97 | 29.31 | 49.21 |
| **CD3+NKG2D^+^** | 38.74 | 46.8 | 52.05 | 39.45 | 32.13 | 59.98 | 25.98 | 23.50 | 46.93 | 58.36 |
| **CD3+CD62L^+^** | 41.19 | 40.78 | 33.94 | 15.66 | 32.97 | 27.31 | 65.36 | 25.10 | 26.65 | 10.72 |
| **CD3+NKp30^+^** | 3.39 | 12.68 | 1.77 | 4.31 | 4.59 | 20.13 | 3.40 | 12.40 | 24.62 | 38.52 |
| **CD3+NKp46^+^** | 0.22 | 0.25 | 0.40 | 1.00 | 0.10 | 0.23 | 0.34 | 0.36 | 0.40 | 0.36 |

CD57^+^ and KLRG-1^+^ cell population was higher in MAD derived CD3+ cells compared with the respected controls.

Patients: MAD1 (Female, 3 y); MAD2 (Male, 5 y), MAD3 (Male, 7 y)
Controls: C1 (Female, 3 y); C2 (Male, 5 y); C3 (Male, 7 y); C4 (Male, 1 y), C5 (Female, 65 y); C6 (Male, 65 y); C7 (Male, 65 y)

**Table S5: Sub-set of NK and B cells.**

| **Cell names** | **markers** | **C1** | **MAD1** | **C2** | **MAD2** | **C3** | **MAD3** | **C4** | **HGPS** | **C5** | **C6** | **C4** |
| --- | --- | --- | --- | --- | --- | --- | --- | --- | --- | --- | --- | --- |
| **NKT cells** | CD3+CD56+ | 0.63 | 1.66 | 1.19 | 1.53 | 1.15 | 1.72 | 2.52 | 0.09 | 4.83 | 4.83 | 13.35 |
| **NK cells** | CD3-CD56+ | 4.43 | 7.97 | 5.28 | 8.69 | 20.82 | 19.75 | 28.46 | 9.54 | 44.24 | 44.24 | 26.99 |
| **NK effector cells** | (CD3-CD56+)CD16^+^ | 94.21 | 93.95 | 94.33 | 91.50 | 98.14 | 97.66 | 95.54 | 96.42 | 97.23 | 97.23 | 96.48 |
| **Cytotoxic NK cells** | (CD3-CD56+)  CD16+CD57+ | 45.04 | 27.65 | 23.40 | 58.39 | 42.82 | 57.25 | 74.25 | 7.33 | 71.87 | 71.87 | 73.88 |
| **B Cells** | CD3-CD19+ | 27.48 | 19.91 | 21.48 | 27.05 | 12.97 | 11.86 | 17.11 | 20.45 | 5.09 | 5.09 | 11.71 |
| **Activated B cells** | (CD3-CD19+)CD25+ | 16.43 | 28.38 | 20.15 | 8.98 | 29.80 | 32.56 | 10.46 | 2.27 | 15.48 | 15.48 | 5.99 |

Patients: MAD1 (Female, 3 y); MAD2 (Male, 5 y), MAD3 (Male, 7 y)
Controls: C1 (Female, 3 y); C2 (Male, 5 y); C3 (Male, 7 y); C4 (Male, 1 y), C5 (Female, 65 y); C6 (Male, 65 y); C7 (Male, 65 y)

**Table S6: Senescent CD3^-^ NK population’s subset**

| **Phenotype (%)** | **C1** | **MAD1** | **C2** | **MAD2** | **C3** | **MAD3** | **HGPS** | **C5** | **C6** | **C7** |
| --- | --- | --- | --- | --- | --- | --- | --- | --- | --- | --- |
| **CD56Bright** | 5.70 | 2.94 | 9.97 | 0.72 | 6.37 | 1.20 | 5.26 | 1.74 | 0.46 | 0.59 |
| **CD56dim** | 94.3 | 97.06 | 90.03 | 99.28 | 93.63 | 98.80 | 94.74 | 98.26 | 99.54 | 99.41 |
| **CD57+** | 34.21 | 30.57 | 21.70 | 59.21 | 35.11 | 50.55 | 18.89 | 60.16 | 71.25 | 68.93 |
| **NKG2A+** | 17.54 | 20.95 | 45.66 | 16.99 | 14.87 | 11.86 | 17.56 | 32.64 | 22.71 | 19.73 |
| **NKG2D+** | 69.30 | 86.23 | 88.42 | 66.73 | 80.52 | 93.55 | 84.42 | 70.48 | 65.93 | 90.13 |
| **CD62L+** | 0.88 | 1.42 | 3.05 | 1.61 | 2.12 | 1.20 | 1.44 | 0.77 | 0.60 | 0.49 |
| **NKp30+** | 43.42 | 37.35 | 40.03 | 26.30 | 28.04 | 43.22 | 64.29 | 65.17 | 58.84 | 56.45 |
| **NKp46+** | 11.40 | 17.91 | 18.49 | 13.60 | 8.51 | 7.38 | 20.33 | 9.19 | 5.21 | 4.93 |
| **CD3+** | 72.45 | 68.69 | 76.99 | 55.05 | 64.80 | 64.10 | 60.50 | 57.87 | 42.62 | 59.52 |
| **CD3-CD56+** | 4.05 | 7.69 | 5.82 | 9.34 | 15.82 | 22.13 | 16.93 | 9.15 | 37.00 | 26.32 |

Patients: MAD1 (Female, 3 y); MAD2 (Male, 5 y), MAD3 (Male, 7 y)
Controls: C1 (Female, 3 y); C2 (Male, 5 y); C3 (Male, 7 y); C4 (Male, 1 y), C5 (Female, 65 y); C6 (Male, 65 y); C7 (Male, 65 y)

**Table S7: T cell phenotypes after cryopreservation of PBMC culture**

| **Markers %** | **C1** | **MAD1** | **C2** | **MAD2** | **C3** | **MAD3** | **C4** | **HGPS** | **C5** | **C6** |
| --- | --- | --- | --- | --- | --- | --- | --- | --- | --- | --- |
| **CD3+** | --- | 99.43 | 99.40 | 98.19 | 99.03 | 99.50 | 97.71 | 99.00 | 99.15 | 98.02 |
| **CD3+CD4+** | --- | 14.86 | 13.97 | 40.81 | 13.97 | 9.58 | 46.85 | 22.26 | 8.84 | 27.58 |
| **CD3+CD8+** | --- | 68.76 | 44.63 | 43.81 | 44.63 | 71.88 | 49.12 | 70.54 | 88.97 | 69.26 |
| **CD4+CD25+** | --- | 4.46 | 4.87 | 21.57 | 4.87 | 2.74 | 8.16 | 5.84 | 2.33 | 4.93 |
| **CD3+CD25+** | --- | 9.46 | 21.55 | 30.22 | 21.55 | 18.88 | 21.95 | 9.91 | 6.34 | 14.31 |

Cells were revived from liquid nitrogen and cultured for three days before confirmation of T lymphocytes. The control group (C1) population was less and was not suited to carry the flowcytometry experiment.

Patients: MAD1 (Female, 3 y); MAD2 (Male, 5 y), MAD3 (Male, 7 y)
Controls: C1 (Female, 3 y); C2 (Male, 5 y); C3 (Male, 7 y); C4 (Male, 1 y), C5 (Female, 65 y); C6 (Male, 65 y); C7 (Male, 65 y)

**Table S8: Levels of Cytokines secreted by cultured PBMC cells.**

| **pg/mL** | **C1** | **MAD1** | **C2** | **MAD2** | **C3** | **MAD3** | **HGPS1** | **C4** | **C5** | **C6** |
| --- | --- | --- | --- | --- | --- | --- | --- | --- | --- | --- |
| **IL-5** | -- | 342.85 | 751.24 | 1691.60 | 524.68 | 525.21 | 13.48 | 150.82 | 138.05 | 522.77 |
| **IL-13** | -- | 198.98 | 431.51 | 863.83 | 227.72 | 250.18 | 16.11 | 83.73 | 17.14 | 123.51 |
| **IL-2** | -- | 5.44 | 3.01 | 3.61 | 6.43 | 5.94 | 13.91 | 2.81 | 9.3 | 4.14 |
| **IL-6** | -- | <0.41 | <0.41 | <0.41 | <0.41 | <0.41 | <0.41 | <0.41 | <0.41 | <0.41 |
| **IL-9** | -- | <0.22 | <0.22 | 1.06 | 1.18 | <0.22 | <0.22 | <0.22 | <0.22 | <0.22 |
| **IL-10** | -- | 1.61 | <0.65 | 4.02 | <0.65 | 1.7 | <0.65 | <0.65 | <0.65 | <0.65 |
| **IFN-γ** | -- | 246.71 | 339.78 | 234.75 | 106.02 | 125.73 | 339.78 | 101.25 | 21.52 | 38.93 |
| **TNF-ɑ** | -- | 1.28 | <1.15 | <1.15 | <1.15 | <1.15 | <1.15 | <1.15 | <1.15 | <1.15 |
| **IL-17A** | -- | <0.96 | <0.96 | <0.96 | <0.96 | <0.96 | <0.96 | <0.96 | <0.96 | <0.96 |
| **IL-17F** | -- | <0.32 | <0.32 | 1.94 | 1.18 | <0.32 | 1.18 | <0.32 | <0.32 | <0.32 |
| **IL-4** | -- | 0.93 | <0.72 | 1.28 | <0.72 | 0.75 | <0.72 | <0.72 | <0.72 | <0.72 |
| **IL-21** | -- | <12.05 | <12.05 | <12.05 | <12.05 | <12.05 | <12.05 | <12.05 | <12.05 | <12.05 |
| **IL-22** | -- | 13 | 0.96 | 40.37 | 68.48 | 12.85 | 0.35 | 0.96 | 0.96 | 0.97 |

Cells were revived from liquid nitrogen and cultured for three days before quantifying the cytokine levels.

Patients: MAD1 (Female, 3 y); MAD2 (Male, 5 y), MAD3 (Male, 7 y)
Controls: C1 (Female, 3 y); C2 (Male, 5 y); C3 (Male, 7 y); C4 (Male, 1 y), C5 (Female, 65 y); C6 (Male, 65 y); C7 (Male, 65 y)

**Table S9: Details of fluorescence conjugated antibodies used in flowcytometry**

| **Specificity and conjugate** | **Dilution** | **Catalog number** | **Supplier** |
| --- | --- | --- | --- |
| APC Mouse Anti-Human CD34 | 1:200 | 560940 | BD Biosciences (Newyork, USA) |
| V450 Mouse Anti-Human CD45 |  | 560368 |  |
| PE Mouse Anti-Human CD73 |  | 561014 |  |
| FITC Mouse Anti-Human CD90 |  | 555595 |  |
| PerCP-Cy5.5 Mouse Anti-Human CD105 |  | 560819 |  |
| Pacific Blue^TM^ Anti-human CD57 Antibody |  | 359608 | BioLegend (San Diego, CA, USA) |
| PE Anti-human CD3 |  | 300308 |  |
| PE Anti-human KLRG1 |  | 36770 |  |
| APC Anti-human CD62L |  | 336012 |  |
| APC Anti-human CD337/NKp30 |  | 325210 |  |
| APC Anti-human CD335/NKp46 |  | 33191 |  |
| APC Anti-human CD57 |  | 322313 |  |
| APC Anti-human CD56 |  | 362503 |  |
| APC Anti-human CD4 |  | 357408 |  |
| PE Anti-human CD8 |  | 344705 |  |
| FITC Anti-human CD25 |  | 302604 |  |
| APC Anti-human TNF-ɑ |  | 502912 |  |
| PE Anti-human IFNγ |  | 506506 |  |
| Anti-human NKG2A/CD159a |  | FAB1059S-100UG | R&D Systems (Minneapolis, MN, USA) |
| Anti-NKG2D/CD314 |  | MAB139-100 |  |

**Table S10: Details of primary antibodies used for western blot analysis**

| **Specificity** | **Dilution** | **Catalog no.** | **Supplier** |
| --- | --- | --- | --- |
| ɑ-tubulin (mouse) | 1:4000 | ab7291 | Abcam (Cambridge, UK) |
| Emerin (rabbit) | 1:1000 | ab156871 |  |
| Histone H4 (rabbit) | 1:1000 | ab177840 |  |
| Histone H3 (acetyl K18) (rabbit) | 1:1000 | ab1191 |  |
| Histone H3 (acetyl K36) (rabbit) | 1:1000 | ab177179 |  |
| Histone H3 (mono methyl K36) (rabbit) | 1:10000 | ab176920 |  |
| Histone H3 (tri-methyl K9) (rabbit) | 1:10000 | ab8898 |  |
| Histone H3 (tri-methyl K4) (rabbit) | 1:10000 | ab8580 |  |
| SUV420h1 (rabbit) | 1:1000 | ab118659 |  |
| GAPDH (mouse) | 1:2000 | AF5009 | Beyotime Institute of Biotechnology (Haimen, China) |
| LaminA/C (mouse) | 1:1000 | 4777S | Cell Signaling Technology (Danvers, MA, USA) |
| LaminB1 (rabbit) | 1:1000 | 17416S |  |
| H4K36me2 (rabbit) | 1:1000 | 2901S |  |
| H4K36me3 (rabbit) | 1:1000 | 4909S |  |
| Tri-Methyl-Histone H3 (Lys27) (rabbit) | 1:1000 | 9763S |  |
| SIRT6 (rabbit) | 1:1000 | 2590S |  |
| TBK1/NAK (rabbit) | 1:1000 | 3504T |  |
| SIRT7 (rabbit) | 1:1000 | 5360S |  |
| Stat3(Tyr705) (rabbit) | 1:1000 | 9145S |  |
| Phospho-Stat3(Ser727) (rabbit) | 1:1000 | 49081S |  |
| STAT3 (mouse) | 1:1000 | 9139S |  |
| SET7/9 (rabbit) | 1:1000 | 2813S |  |
| MTX2 (rabbit) | 1:1000 | 11610-1-AP | ProteinTech Group, Inc. (Rosemont, IL, USA) |
| MTX1 (rabbit) | 1:1000 | 15529-1-AP |  |
| DRP1 (rabbit) | 1:1000 | 12957-1-AP |  |
| MFN2 (rabbit) | 1:1000 | 12186-1-AP |  |
| LaminB2 (rabbit) | 1:1000 | 10895-1-AP |  |
| Histone 3 (rabbit) | 1:2000 | 17168-1-AP |  |
| cGAS (rabbit) | 1:1000 | 26416-1-AP |  |
| TMEM173/Sting (rabbit) | 1:1000 | 19851-1-AP |  |
| AIM2 (rabbit) | 1:1000 | 20590-1-AP |  |
| NLRP3 (rabbit) | 1:1000 | 19771-1-AP |  |
| IL6 (mouse) | 1:1000 | 21865-1-AP |  |
| COXIV (rabbit) | 1:1000 | A11631 | ABclonal (Woburn, MA, USA) |
| VDAC1 (rabbit) | 1:1000 | A19707 |  |
| Phospho-TBK1/NAK-S172 (rabbit) | 1:1000 | AP1026 |  |
| p62/SQSTM1 (rabbit) | 1:1000 | P0067 | Sigma-Aldrich; Merck KGaA (Darmstadt, Germany) |
| LC3B (rabbit) | 1:1000 | L7543 |  |
| Histone H3ac (pan-acetyl) (rabbit) | 1:1000 | 39139 | Active Motif, Inc. (Carlsbad, CA, USA) |
| Histone H3R17me2a (rabbit) | 1:10000 | 39709 | Thermo Fisher Scientific, Inc. (Waltham, MA, USA) |
| Histone H3R2me2S (rabbit) | 1:10000 | ABE460 | Millipore; Merck KGaA (Darmstadt, Germany) |
| SIRT1 (rabbit) | 1:1000 | SC-15404 | Santa Cruz Biotechnology, Inc. (Santa Cruz, CA, USA) |
| SUV420h2 (rabbit) | 1:1000 | NBP2-94300 | Novus Biologicals (Littleton, CO, USA) |

**Table S11: Details of primary antibodies used for immunofluorescence**

| **Specificity** | **Dilution** | **Catalog no.** | **Supplier** |
| --- | --- | --- | --- |
| Lamin A/C (mouse) | 1:300 | 4777S | Cell Signaling Technology (Danvers, MA, USA) |
| LaminB1 (rabbit) | 1:300 | 17416S |  |
| γ-H2AX (rabbit) | 1:300 | 7631S |  |
| Phospho-Stat3(Tyr705) (rabbit) | 1:300 | 9145S |  |
| Phospho-Stat3(Ser727) (rabbit) | 1:300 | 49081S |  |
| OCT4,TRA160, SOX2, SSEA4 | 1:300 | ab109884 | Abcam (Cambridge, UK) |
| Emerin (rabbit) | 1:300 | ab156871 |  |
| LaminB2 (rabbit) | 1:300 | 10895-1-AP | ProteinTech Group, Inc. (Rosemont, IL, USA |

**Table S12: List of Primers**

| Name | Sequences (5’ to 3’) |
| --- | --- |
| LMNA-Forward | AATGATCGCTTGGCGGTCTAC |
| LMNA-Reverse | CACCTCTTCAGACTCGGTGAT |
| LMNB1-Forward | GAAAAAGACAACTCTCGTCGCA |
| LMNB1-Reverse | GTAAGCACTGATTTCCATGTCCA |
| LMNB2-Forward | GTCCTGGATGAGACGGCTC |
| LMNB2-Reverse | GCGCTCTTGTTGACCTCGT |
| IL1β-Forward | ATGATGGCTTATTACAGTGGCAA |
| IL1β-Reverse | GTCGGAGATTCGTAGCTGGA |
| IL6-Forward | GCCACTCACCTCTTCAGAAC |
| IL6-Reverse | GCAAGTCTCCTCATTGAATCCA |
| IL10-Forward | GACTTTAAGGGTTACCTGGGTTG |
| IL10-Reverse | TCACATGCGCCTTGATGTCTG |
| IL8-Forward | CGGAAGGAACCATCTCACTGT |
| IL8-Reverse | GGTCCACTCTCAATCACTCTCA |
| IL18-Forward | TCTTCATTGACCAAGGAAATCGG |
| IL18-Reverse | TCCGGGGTGCATTATCTCTAC |
| IL23-Forward | CTCAGGGACAACAGTCAGTTC |
| IL23-Reverse | ACAGGGCTATCAGGGAGCA |
| TGFβ1-Forward | GGCCAGATCCTGTCCAAGC |
| TGFβ1-Reverse | GTGGGTTTCCACCATTAGCAC |
| IFNβ-Forward | TCTCCTCCAAATTGCTCTCC |
| IFNβ-Reverse | CTCCCATTCAATTGCCACAG |
| LINE1-Forward | TCAGGTTGACAGCAGACTTATC |
| LINE1-Reverse | CCTGGCGTGCTAAGGTATATT |
| TFAM-Forward | GTTGGAGGGAACTTCCTGATT |
| TFAM-Reverse | CGTTATAAGCTGAACGAGGTCT |
| AIM2-Forward | TGGCAAAACGTCTTCAGGAGG |
| AIM2-Reverse | AGCTTGACTTAGTGGCTTTGG |
| NLRP3-Forward | GATCTTCGCTGCGATCAACAG |
| NLRP3-Reverse | CGTGCATTATCTGAACCCCAC |
| IFNα-Forward | GCCATCTCTGTCCTCCATGA |
| IFNα-Reverse | GCTGGTAGAGTTCGGTGCAG |
| IFNβ-Forward | GCCGCATTGACCATCTATGA |
| IFNβ-Reverse | AGTCTCATTCCAGCCAGTGCT |
| TNFα-Forward | TGGAGAGTGAACCGACATGG |
| TNFα -Reverse | CTCTCAGCTCCACGCCATT |
| cGAS-Forward | CTCCACGAAGCCAAGACCTC |
| cGAS-Reverse | GCGGCTGAGCTTCAACTTCT |
| STING-Forward | CCTGTTGCTGCTGTCCATCT |
| STING-Reverse | ATGTTCAGTGCCTGCGAGAG |
| TBK1-Forward | GGAAGCGGCAGAGTTAGGTG |
| TBK1-Reverse | TCGGATGAGTGCCTTCTTGA |
| IRF3-Forward | AGAGGCTCGTGATGGTCAAG |
| IRF3-Reverse | AGGTCCACAGTATTCTCCAGG |
| p53-Forward | TGAGGTTGGCTCTGACTGTA |
| P53-Reverse | GTGTGATGATGGTGAGGATGG |
| p16-Forward | CTCTGAGAAACCTCGGGAAAC |
| p16-Reverse | ATGAAAACTACGAAAGCGGG |
| p21-Forward | CTCTACATCTTCTGCCTTAGTCTCA |
| p21-Reverse | ACCTCTCATTCAACCGCCTA |
| MTX1-Forward | GTGCTGACCTATGCCAGATTTA |
| MTX1-Reverse | CCACGTAGTTCTTGGTGTCTATC |
| MTX2-Forward | ACTGGGAACACAACCGTATTT |
| MTX2-Reverse | CTCTATGACAGCCTGCCTTTAC |
| SAMM50-Forward | GAGCCTGAAGCTAAACAGGAA |
| SAMM50-Reverse | TCTGCACGACCAAGAAGATTAG |
| VDAC2-Forward | GGACCTTGGAGACCAAATACA |
| VDAC2-Reverse | CCAGCCCTCATAACCAAAGA |
| CHCHD3-Forward | GTTCGAAGTCTCAGCGGTATT |
| CHCHD3-Reverse | CAGTCTAGCCAGCTGTTCTTT |
| TOMM22-Forward | CCTTTCACCAAATTGCTCCTAAC |
| TOMM22-Reverse | GCCTCCCTCCTCTCATACATA |
| OPA1-Forward | GAGGACAGCTTGAGGGTTATTC |
| OPA1-Reverse | GTTCTTCCGGACTGTGGTTATT |
| DRP1-Forward | GGTGAACCCGTGGATGATAAA |
| DRP1-Reverse | GACGAGGACCAGTAGCATTTC |
| MFN1-Forward | CAGTGGGAAGAGCTCTGTTATC |
| MFN1-Reverse | TGTGCCTGGACTGTCTACTA |
| NDUFS1-Forward | ACCTGGACTTGGGATGAAATAC |
| NDUFS1-Reverse | GAGCCTTCTGGGAGATGATTAG |
| MFN2-Forward | CATGCAGCAGGACATGATAGA |
| MFN2-Reverse | ATAGACGTAGAGGAGGCCATAG |
| DRP1-Forward | GGTGAACCCGTGGATGATAAA |
| DRP1-Reverse | GACGAGGACCAGTAGCATTTC |
| GAPDH-Forward | TCGGAGTCAACGGATTTGGT |
| GAPDH-Reverse | TTGCCATGGGTGGAATCATA |

**Table S13: crRNA and Donor template sequences.**

|  | Sequence (5’ to 3’) | bp |
| --- | --- | --- |
| Alt-R CRISPR-Cas9 crRNA | CTGCGGGAACAGCCTG**T**GTA | 20 |
| Alt-R HDR Donor Oligo (ssDNA) | TGTGGAAGGCACAGAACACCTGGGGCTGCGGGAACAGCCTGCGTACGGCTCTCATCAACTCCACTGGGGAAGTAAGTAGGCCTG | 84 |
